# Barriers to Antimicrobial Resistance Gene Exchange in Methicillin Resistant *Staphylococcus aureus* Cluster into Transfer Islands

**DOI:** 10.64898/2025.12.19.694131

**Authors:** Jacob Wildfire, Mais Maree, Sophie R. Mallinson, Adam A. Witney, Gwenan M. Knight, Jodi A. Lindsay

## Abstract

Horizontal gene transfer (HGT) via generalised transduction is a major driver of antimicrobial resistance (AMR) in *Staphylococcus aureus*, yet the genetic barriers regulating phage-mediated transfer remain poorly defined. Using a co-culture gene transfer rate assay (COGTRA) that quantifies phage-dependent resistance gene exchange under competitive, antibiotic-free conditions, we screened 1,920 mutants from the Nebraska Transposon Mutant Library for elevated transfer. We identified 32 validated high-transfer mutants that act as HGT barriers, 66% of which clustered within two chromosomal regions that we term *S. aureus* Transfer Islands (SauTI1 and SauTI2). SauTI loci restricted generalised transduction bidirectionally, and SauTI1 genes were widespread across *S. aureus* genomes yet showed lineage-specific variation. SauTI1 encodes an SMC2–Cbf1 condensin–nuclease module whose disruption markedly increases generalised transduction and plasmid transfer, resembling Wadjet-like restriction systems. Together, these findings identify clustered defence loci that restrict HGT in *S. aureus* and identify SauTIs as key regulators of AMR evolution.

## Introduction

*Staphylococcus aureus* is a common commensal of humans and animals. As an opportunistic pathogen, it can cause a wide variety of infections ranging from mild to life-threatening (1). Importantly, Methicillin resistant *S. aureus* (MRSA) is the leading cause of antimicrobial resistance (AMR)-associated fatalities worldwide, accounting for over 100,000 deaths annually (2, 3).

Despite its remarkably conserved core genome, *S. aureus* exhibits high adaptability to diverse environments, mainly through the acquisition of mobile genetic elements (MGEs) that encode resistance, virulence, and host adaptation genes (3, 6, 7). The high prevalence of diverse MGEs, including plasmids, transposons, and prophages, in *S. aureus* genomes, highlights the central role of horizontal gene transfer (HGT) in shaping its evolution (4). Among the major HGT mechanisms, including conjugation and natural transformation, generalised transduction is considered the dominant route (5). In this mechanism, bacteriophage (henceforth referred to as “phage”) package and transfer bacterial DNA from anywhere in the genome from one host cell to another (4, 8, 9).

While HGT occurs extensively during *S. aureus* colonisation (6), it remains a notoriously difficult bacteria to genetically modify in *in vitro* conditions (7, 8). This contrasts with the significant generalised transduction efficiency of antimicrobial resistance genes (ARGs) during *in vivo* co-colonisation of gnotobiotic piglet skin, a model of human colonisation, even in the absence of antibiotic selection (6). Furthermore, many MGEs seem to be restricted by lineage, particularly the functionally cryptic *S. aureus* genomic islands νSAα and νSAβ (9–11). The discovery of the Sau1 restriction modification (R-M) *hsdM* and *hsdS* genes, each on a genomic island and correlating strongly with lineage, was initially thought to be responsible for *S. aureus* HGT control (12). However, knockout of the core restriction gene (*hsdR*) is not sufficient to permit very high transfer rates (13). Although a second R-M system, SauUS1, that blocks transfer from *Escherichia coli* has been discovered, how *S. aureus* governs HGT remains unclear (14).

One possible explanation may lie in tightly controlled and as-yet undiscovered transduction activation pathways. We have previously demonstrated that sub-inhibitory concentrations of antibiotics increase the rate of generalised transduction, and that this increase is not correlated with prophage induction, but rather the rate of non-prophage DNA packaging (15). This raises the possibility that HGT is induced via an unknown signalling pathway. Equally, there may be undiscovered genetically encoded barrier systems. New HGT barrier systems have been recently revealed in a myriad of other bacterial species via the bioinformatic screening of thousands of genomes by identifying clusters of putative genes and subsequently validating their phenotype (16, 17). These systems act to restrict incoming foreign DNA at various stages of transfer (Supplementary Fig. 1). To date, no new barrier genes have been described in *S. aureus* using this approach. Given the difficulty of genetically modifying *S. aureus*, it is plausible that novel systems remain undiscovered. Examples of new systems in other bacteria include the recently described Wadjet system, a condensin complex that recognises plasmids by completely extruding looped DNA through its ring-like structure before cleaving them via a nuclease (18, 19). Here, we sought to identify genes that control HGT in this species to better define gene transfer dynamics. Such insights will help overcome current genetic modification limitations and shed light on how successful, increasingly resistant *S. aureus* strains emerge and dominate.

To achieve this, we conducted a phenotypical screen of the well characterised Nebraska Transposon Mutant Library (NTML) of 1,920 *S. aureus* gene disruptions in the clinical MRSA JE2 strain (20). Using our established co-culture gene transfer rate assay (COGTRA) (21), each of the individual transposon mutants harbouring the erythromycin ARG, *ermB*, was co-cultured with the ‘fixed strain’, an *ermB* negative, kanamycin resistant mutant in the JE2 background carrying the pT181 tetracycline resistance plasmid (Fig. 1a). The rate of ARG transfer under competitive pressure is measured by the concentration of double resistant progeny (DRP) to erythromycin and tetracycline. We also explored condensin mutants due to their knockout gene’s chromosomal position and homology with the Wadjet barrier system.

**Figure 1.**
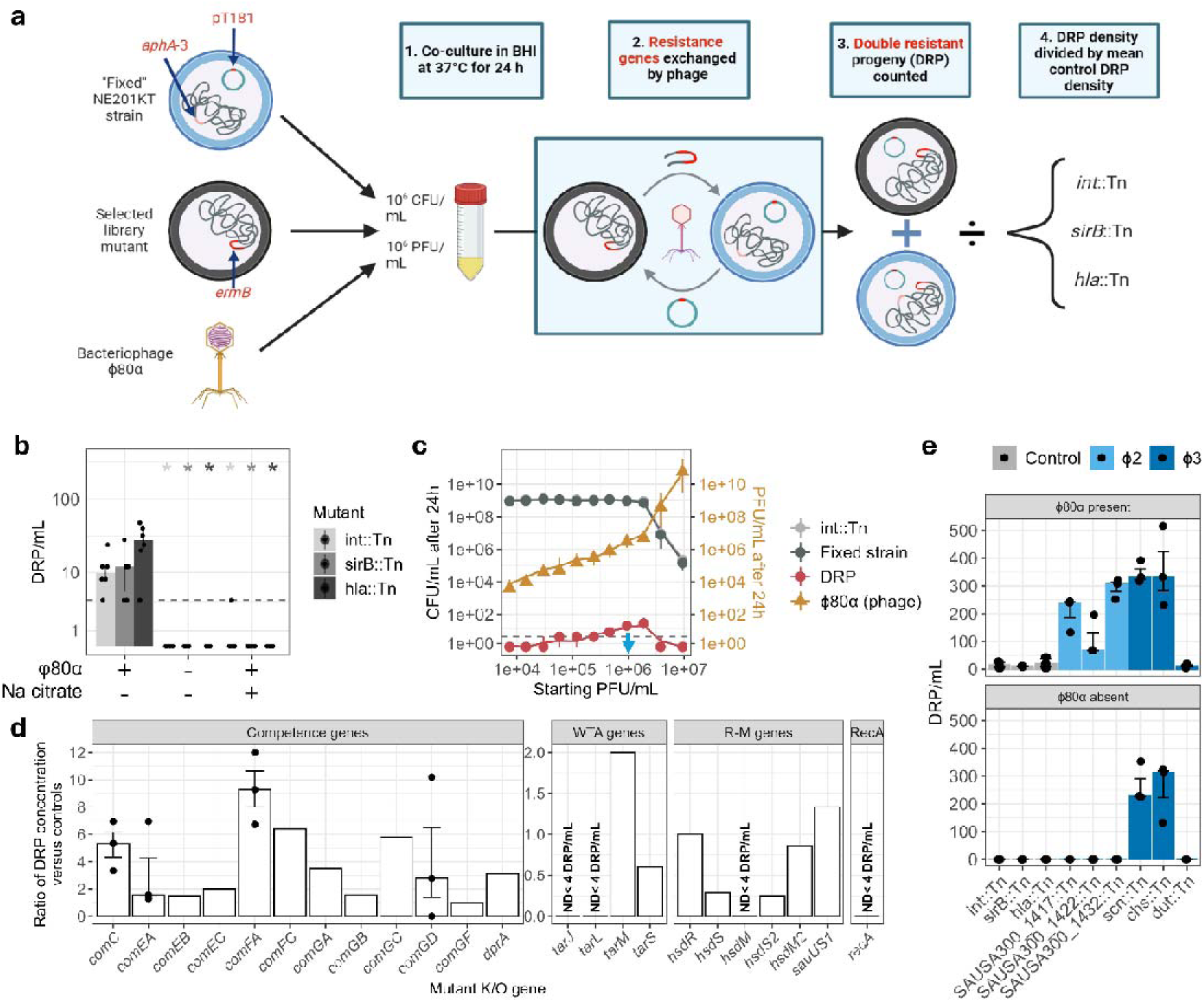
Generalised transduction via φ80α is the primary mechanism of HGT for non-prophage loci in the COGTRA. (a) Diagram of the COGTRA screening methodology. A selected library mutant, which has a known gene interruption caused by a transposon containing an erythromycin resistance gene *(ermB),* is used to inoculate brain heart infusion media (BHI) to a concentration of 10^6^ CFU/mL. NE201KT, or the “fixed strain”, which contains a kanamycin resistance gene (*aphA*-3) and the pT181 tetracycline resistance plasmid, is added at the same concentration, along with generalised transducing phage φ80α to a concentration of 10^6^ PFU/mL. 1) The co-culture is then incubated at 37°C for 24 hours (h). 2) During incubation, antimicrobial resistance genes, indicated on the chromosome or plasmid with red or pink, are exchanged via φ80α. 3) After 24 hours, the rate of gene transfer is measured by calculating the concentration of double resistant progeny (DRP), which are resistant to both erythromycin and tetracycline. 4) To provide standardisation, the DRP density of the selected mutant is divided by the mean DRP density of three control mutants grown in respective, individual co-cultures with the fixed strain at the same time as the selected mutant, *int*::Tn, *sirB*::Tn *and hla*::Tn, which contain knockouts in genes which do not affect the rate of generalised transduction. Created with Biorender.com. (b) Control mutant *(int*::Tn, *sirB*::Tn and *hla*::Tn) COGTRA in the presence or absence of φ80α or 20 mM Na citrate. Points represent the DRP concentration (y-axis) of each replicate (n = 6) after 24 hours, with bar values representing the median. Error bars are the IQR. DRP detection limit is indicated by dashed line. Significant difference between each mutant condition and their respective mutant control, with φ80α and without Na citrate, was determined via Wilcoxon rank-sum test with Bonferroni correction; no asterisk > 0.05, * < 0.05, with colour denoting the NTML control mutant. (c) Co-cultures varied by starting φ80α concentration (x-axis). After 24 hours of incubation, parent and DRP strain concentrations (CFU/mL, left y-axis), as well as φ80α concentrations (PFU/mL, right y-axis), were determined for each independent co-culture. The chosen optimal starting phage concentration is indicated by the blue arrow. Values are medians (n = 9); error bars represent the IQR. DRP detection limit is indicated by dashed line. Note that the control mutant (*int*::Tn) time series lies behind the fixed strain. (d) COGTRA of mutants of genes that play a role in *S. aureus* HGT. For most mutants, the DRP concentration of a single replicate (n = 1) after 24 hours was divided by the mean DRP concentration of three control mutants, which is represented by a respective bar. Mutants *comC*::Tn, *comEA*::Tn, *comFA*::Tn and *comGD*::Tn, however, were tested in triplicate (n=3), with points representing each independent replicate, the bar values representing the median and error bars the IQR. Mutants with co-culture below the detection limit are indicated via labelling. (e) COGTRA of mutants with disrupted genes located on the JE2 prophage and control mutants, in the presence or absence of φ80α, demonstrates that φ80α is required for the transfer of φ2. Points represent the DRP concentration of each replicate (n = 3) after 24 hours, with bar values representing the median. Bar colour represents the prophage. Error bars are the IQR. (b-e) The limit of detection was 4 DRP/mL. Values below this were given a concentration of 0 DRP/mL. n represents the number of independent replicates.

This screen identified 32 genes whose disruption led to significantly increased gene transfer, suggesting that they function either as transfer barriers or transduction-activation genes. We found that many of these genes cluster in two genomic regions conserved across most of the strains that we analysed; we have termed these the *Staphylococcus aureus* Transfer Islands (SauTIs). SauTI1 encodes *hsdS* as well as previously unrecognised accessory condensin and Wadjet-like *jetCD* homologs that functionally controls gene transfer between *S. aureus* strains, that we have called SMC2-Cbf1.

## Results

### Generalised transduction is the primary mechanism of resistance gene transfer in co-culture

The COGTRA assay measures HGT by co-culturing two parent strains, each with a unique ARG, in Brain Heart Infusion (BHI) broth for 24 hours and counting DRPs generated following selection on ARG-specific antibiotics (Fig. 1a) (21). JE2, the background of the mutant library, lacks conjugative genes. JE2 strains also fail to exchange DNA unless exogenous generalised transducing bacteriophage φ80α is added. This transfer is blocked by citrate which chelates calcium and prevents phage binding as confirmed in three control mutants *int*::Tn, *sirB*::Tn and *hla*::Tn (Fig. 1b). These findings indicate that the observed transfer is phage dependent. Phage concentrations (PFU/mL) positively correlated with DRP concentration up to 2×10^6^ colony forming units per millilitre (CFU/mL), further supporting that φ80α is responsible for ARG transfer (Fig. 1c). However, higher starting concentrations produced a sharp decline in DRP and parent concentrations, likely due to excessive φ80α-mediated bacterial lysis as demonstrated by the large increase in phage concentration (Fig. 1c). A starting concentration of 1×10^6^ plaque forming units per millilitre (PFU/mL) was therefore selected for subsequent library screening.

Natural transformation is another possible HGT mechanism, enabling *S. aureus* to acquire exogenous DNA through the expression of competence-associated genes under specific environmental conditions (22, 23). While addition of DNase prevented gene transfer (Supplementary Fig. 2), deletion of 12 known competence genes did not affect DRP ratios (Fig. 1d), suggesting that DNase may interfere with the exogenous phage (Supplementary Fig. 2). In contrast, biosynthesis genes of wall teichoic acid (WTA), the receptor for *S. aureus* phage (24), were necessary for gene transfer. Mutants *tarJ*::Tn and *tarL*::Tn, WTA biosynthesis pathway knockouts, produced DRPs below the detection limit, whereas mutants of the WTA glycosylation genes *tarM* or *tarS*, demonstrated ratios of 2.00 and 0.60 respectively. This might be expected given the function of WTAs as phage binding targets; their absence would totally inhibit transduction. Whilst in the absence of one *tar* gene the other may provide adequate glycosylation (25).

None of the known Sau1 R-M mutants produced an increased DRP ratio (Fig. 1d). This was expected, because the parent strains share the same JE2 background. As such, HsdR-mediated restriction is unlikely, since transferred DNA would be correctly modified (26). Indeed, an *hsdR*::Tn mutant produced a DRP ratio of 1.00 compared to the control mutants. In contrast, the other Sau1 R-M mutants, *hsdS*::Tn, *hsdM*::Tn, *hsdS2*::Tn and *hsdM2*::Tn, demonstrated decreases in DRP ratios. Decreases are expected as when the DNA is not modified correctly in the donor bacteria, the delivered DNA will be recognised by a functioning HsdSMR complex in the recipient bacteria and digested. However, as *hsdM2*::Tn had a ratio of 0.00, these results also suggest that whilst *hsdM*1 on νSAα is functional, *hsdM*2 on νSAβ is not. Both copies of *hsdS* are functional. The R-M SauUS1 does not impact on gene transfer in this assay. Other genes likely implicated in HGT such as the DNA integration gene, *recA*::Tn, produced a ratio of 0.00, indicating that transfer is inhibited in the absence of functioning homologous recombination.

JE2 carries two prophage in its genome, φ2 and φ3, but they are not sufficient for generalised transduction in this assay. φ2 genes could transfer only in the presence of exogenous φ80α (Fig. 1e). In contrast, φ3 genes could transfer in the absence of φ80α, suggesting the prophage is mobile, but not able to package and deliver non-phage DNA.

### Two independent genomic clusters of transfer barrier gene candidates exist in the chromosome

Following optimisation and validation of the COGTRA, we then screened the transposon library for high transfer mutants (HTM) to identify genes that in the wildtype may control HGT (Supplementary Table 1). The dataset was substantially right skewed, with most mutants demonstrating DRP ratios clustering around 1 and therefore indicating no difference relative to controls, and a subset demonstrating very high ratios indicating a high increase (Supplementary Fig. 3a). This suggests that most gene knockouts have little impact on HGT, whereas a small number have a strong impact. A DRP ratio of 13 was chosen as the cut-off to identify HTMs as this was the 95^th^ percentile; this produced 98 HTMs with ratios of 13 or greater (Supplementary Table 1). Two of these harbour the same gene knockout (SAUSA300_1802::Tn), and so one was excluded, yielding 97 HTMs. Analysing the distribution of these HTMs along the chromosome revealed that many have knockouts in genes opposite the origin of replication, an observation supported by a trend line which shows a general increase in this region (Fig. 2). This is partly explained by a high density of HTM genes within φ2 (8 HTMs) and φ3 (9 HTMs); as we have evidenced that it is possible that these are the result of prophage replication, φ2 and φ3 HTMs were excluded from further interrogation leaving 80 HTM genes. However, 52 of these (65%) reside between the prophage. As demonstrated in the histogram of HTMs per 50 kb, these genes form two separate clusters between, but not next to, the prophage loci. This is supported by statistical analysis, which revealed that the HTM gene loci pattern is non-random (Supplementary Fig. 3b). Together, this suggests the existence of dedicated regions for genes that control HGT within the *S. aureus* chromosome. The larger and more downstream of the two clusters, which we term the *Staphylococcus aureus* Transfer Island 1 (SauTI1), contains 28 of the 80 HTM genes (35%), as well as two Type I R-M genes (*hsdS* and *hsdM*) (Fig. 2). Based simply on the location of the HTM genes, the approximate span of SauTI1 is 1923706-2014500 bp. Interestingly, the left half overlaps with the genomic island νSAβ, which is known to vary according to clonal complex (11), whilst the right half sits within the conserved chromosome. The smaller and more upstream cluster, SauTI2, contains 16 HTM genes (20%), and an approximate locus of 1753253-1846903 bp.

**Figure 2.**
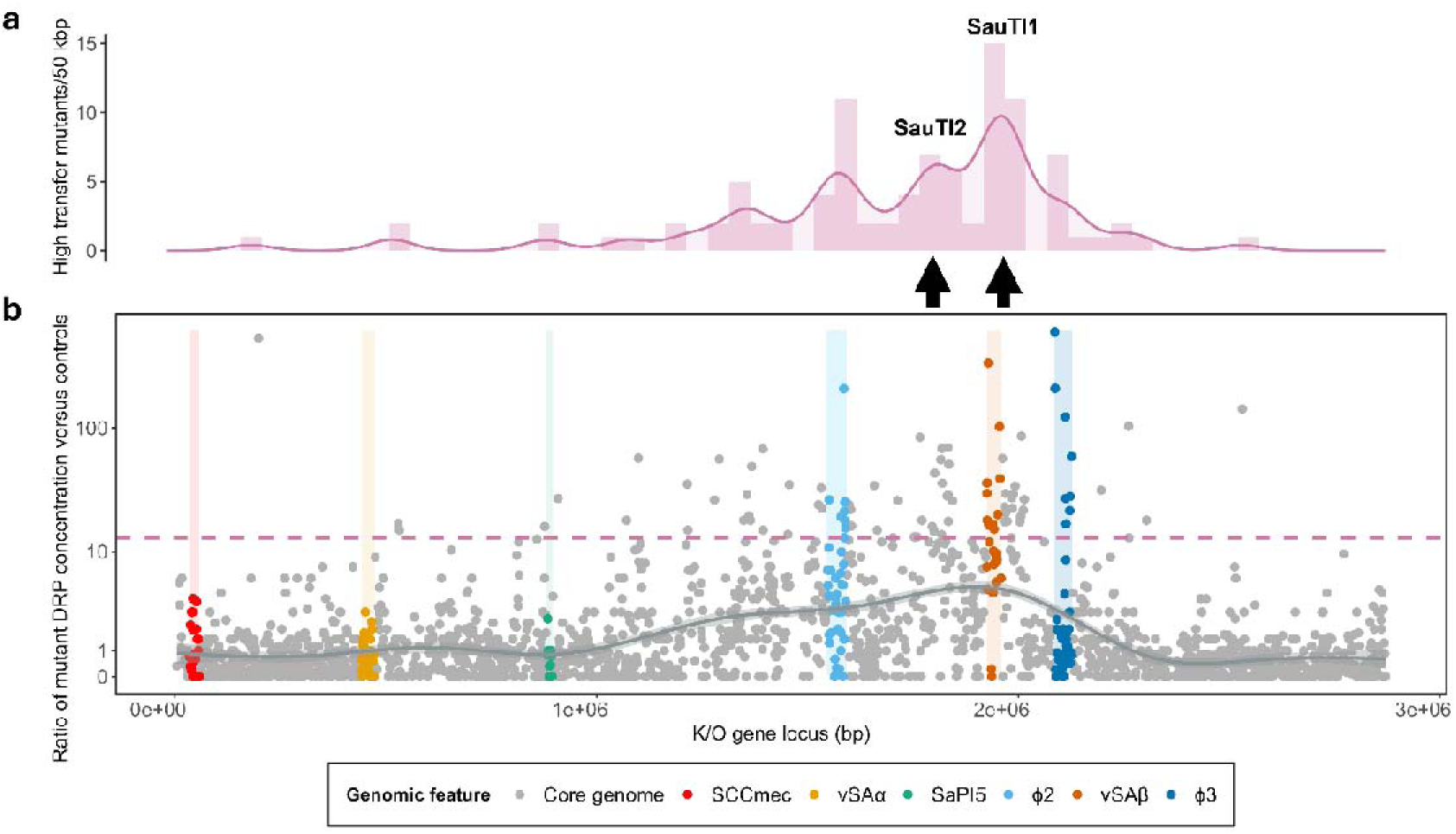
Screen of the entire mutant library showing mutants that demonstrate high frequency transfer clustering between, but not next to, φ2 and φ3. (a) Histogram of high transfer mutants per 50 kb. Trend line (solid pink) plotted via kernel density estimation. The *Staphylococcus aureus* Transfer Islands (SauTI) 1 and 2 are each indicated by a respective black arrow. (b) Each mutant’s DRP concentration was divided by the mean DRP concentration of three control mutants (y-axis) and plotted against the mutant’s knockout (K/O) gene start locus (x-axis). Dots represent each mutant (n = 1). Trend line (grey) plotted via kernel density estimation. MGEs and corresponding genes are indicated by colour (legend bottom). High transfer mutant threshold (≥ ratio of 13) is represented by the pink dashed line.

In addition, only 385 mutants (20% of the library) did not produce DRPs, and these mutants did not show any discernible pattern in their gene loci (Fig. 2). Therefore, most mutants could produce DRPs in co-culture, regardless of knockout gene locus. Those that did not produce DRPs may have knockouts in genes essential for HGT, however, the assay was not designed to accurately detect such mutants, or those that promote HGT. Further analysis is therefore required to verify genes essential to or that promote HGT.

### The screen was highly sensitive for the detection of high transfer

Given that only a single COGTRA repeat was performed for each of the 1,920 mutants, we next sought to evidence that the screen results were reproducible and confirm the transfer phenotype of the HTMs. We therefore repeated the COGTRA methodology in multiplicate on all 80 non-prophage HTMs and a selection of 49 non-prophage low transfer mutants (LTM) that represented the entire chromosome. The results were then used to compare the results of the singular COGTRA screen. Of the 129 total mutants tested by the multiplicate screen, four prospective HTMs produced fewer than three successful replicates and so were not included in the analysis of the singular screen. 89 of the remaining 125 mutants (71.2%) showed a multiplicate DRP ratio decrease, suggesting that the singular COGTRA screen produced generally higher ratio values (Supplementary Fig. 4a). The multiplicate COGTRA, which was used as the gold standard, yielded 32 HTMs and 93 LTMs (Supplementary Table 2 and Supplementary Fig. 4a). All multiplicate HTMs were also high transfer in the singular screen, and so there were no false negatives. The sensitivity of the singular screen was therefore 100% for the detection of HTMs, and as such likely detected all non-prophage HTMs. In contrast, 44 of the 93 LTMs identified by the multiplicate COGTRA were HTMs in the singular screen and were therefore false positives. This indicates that the singular screen was 52.7% specific for the detection of HTMs, and therefore some HTM candidates were incorrectly identified. However, this was resolved by the performance of the multiplicate COGTRA on 76/80 non-prophage HTMs, allowing false positives to be removed, providing a list of “confirmed” HTMs. Despite fewer than three successful repeats, SAUSA300_1747::Tn was included due to its incredibly high DRP ratio. In addition, genotypic analysis of the HTM *kdpA*::Tn revealed that the mutant was a repeat of the HTM SAUSA300_1785::Tn, resulting in its removal; this left a remaining 32 confirmed transfer barrier gene candidates (Supplementary Table 3). Seven of these were putative genes of unknown function and may therefore have new mechanisms for controlling DNA transfer.

To confirm the presence of the SauTI1 and SauTI2 clusters in the multiplicate COGTRA HTMs, we analysed the multiplicate COGTRA and explored the mutant knockout loci. Clustering was still visible within the SauTI loci, with 11 (34.4%) clustered within SauTI1 (1927784-1988370 bp), and 10 (31.3%) clustered within SauTI2 (1801077-1846903 bp) (Supplementary Fig. 4b-c). Statistical analysis provided further evidence that the HTM distribution along the chromosome is not random, further confirming the SauTI existence (Supplementary Fig. 4d).

### Most HTMs show increased ARG transfer out of the mutant or in both directions

The identified candidate genes could control HGT at any step required for successful DNA transfer (Supplementary Fig. 1). Identifying which step would help to characterise how it functions to control DNA transfer. Given that only the fixed strain harbours a kanamycin ARG, we were able to measure the direction of transfer by determining whether DRPs were kanamycin resistant. This was used to determine the rate of transfer in each direction for all 32 confirmed HTMs and 93 LTMs. The concentration of kanamycin sensitive (KanS) and kanamycin resistant (KanR) DRPs were each divided by the overall DRP density of control mutants to provide standardisation. A KanS or KanR ratio of 10 or greater was chosen to highlight strains with a marked elevation relative to the control, as a value of 10 for one ratio indicates 10 times the overall transfer rate of controls in that direction alone. Of the 32 HTMs, 20 HTMs (60.6%) demonstrated an increase in the KanR ratio exclusively. This indicates elevated ARG donation from mutants to the fixed strain, implying increased mobility or packaging of the knockout locus (Supplementary Table 3). In contrast, 2 (6.3%) demonstrated an increase of 10 or greater in the KanS ratio exclusively, indicating increased ability to receive the resistance plasmid. This therefore suggests the wildtype genes are receiver barriers. Finally, a further 9 HTMs (27.3%) demonstrated both KanS and KanR ratios above 10. These HGT increases in both directions could potentially be due to the knocked-out gene being involved in HGT signal transduction. Their knockout could therefore activate generalised transduction in the mutant, and where the knockout is transferred to fixed strain, creating a DRP, generalised transduction could similarly activate increasing plasmid transfer. The genes of the 20 HTMs with increased *ermB* transfer cluster together along with the genes of the 9 mutants with increased transfer in both directions, particularly within SauTI1 and SauTI2 (Supplementary Fig. 5a and Supplementary Table 3).

To investigate whether SauTIs might be mobile, selected mutants spanning SauTI1 were tested and appeared to have a marginally increased KanR ratio, suggesting some mobility (Supplementary Table 3). However, when we transferred in the absence of φ80α, there was no indication that SauTI moves on its own (Supplementary Fig. 5b).

### SauTI1 varies between different clonal complexes but less so within

Given the SauTI’s potential importance in the control of HGT and the overlap of SauTI1 with νSAβ, a feature strongly correlated with lineage (11), we next searched for SauTI1 homologues within other strains. This was to understand how the SauTI1 is distributed within the *S. aureus* population, as well as the degree to which it varies within and between clonal complexes (CC). Homologues were identified in 21 out of 22 well annotated, complete chromosomal sequences from nine clonal complexes (Fig. 3). Comparison of their gene homology revealed substantial conservation in the posterior region of the SauTI1, with the nine HTM knockout gene loci located in the posterior region being present in all isolates (Supplementary Fig. 6). Strains JH1 and JH9 (CC5) differ minorly due to a transposase. In contrast, the anterior region, which overlaps with νSAβ, demonstrates greater variability. Homologues of the two anterior HTM knockout gene loci, SAUSA300_1747 and SAUSA300_1769, appeared in 12 (54.5%) and 17 (77.3%) of isolates, respectively. However, most variation appears to be between different clonal complexes; for example, the anterior regions of CC5, CC22 and CC30 isolates appeared to harbour an enterotoxin cluster, which was replaced by a lantibiotic epidermin cluster in CC1, CC8, CC151 and CC425 (Fig. 3). Isolates within a CC-type generally showed little deviation from each other, except within CC239: JKD6008 showed a 26.0 kb loss of the anterior region which was not shared by 0582/TW20. Two animal isolates, S0385 (CC398) and RF122/ET3-1 (CC151), looked substantially different from the human isolates. The swine isolate S0385 demonstrated a large deletion in the anterior SauTI1 region, whereas bovine isolate RF122/ET3-1 demonstrated a 43.3 kb insertion. In contrast, another bovine isolate LGA251 (CC425) greatly resembled isolates of CC1 and CC8, and avian isolate ED98 (CC5) showed close homology to other members of CC5. This suggests that complexes comprised of strains primarily isolated from animal hosts have more substantial variation.

**Figure 3.**
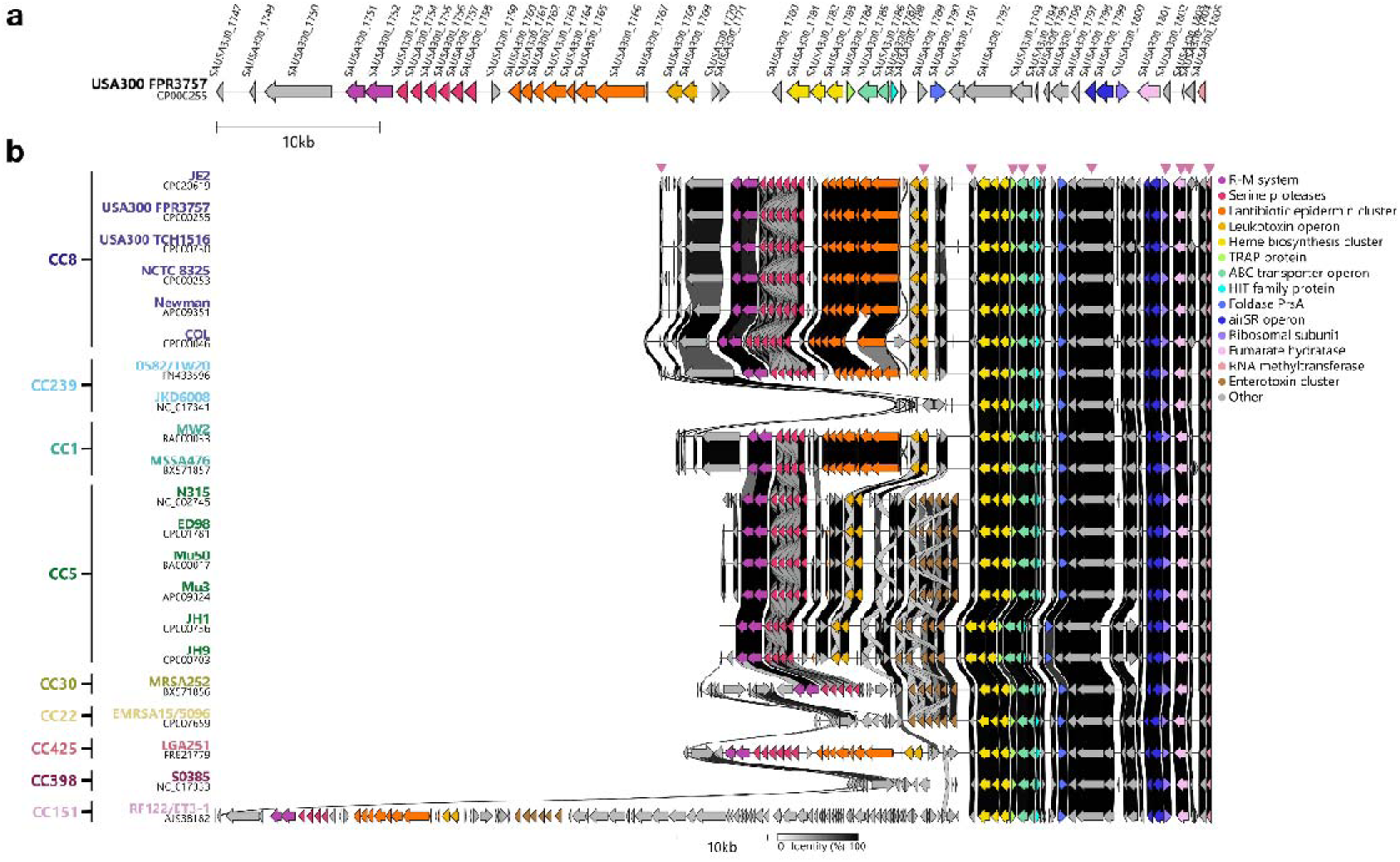
Gene homology between SauTI1 sequences by clonal complex (CC). (a) Schematic map of the USA300_FPR3757 SauTI1. The respective locus is presented at the start of each gene. (b) Comparison of the SauTI1 homologues of 21 isolates. Isolate name and accession number are left of the figure, with respective colour representing the respective CC. HTMs indicated by pink triangles above the first sequence. Connecting bands represent the degree of overall identity between the gene homologues stacked above it (identity bar, bottom). Sequences were sorted into CCs, organised by primary host organism (human, bovine and swine), with CC8, and therefore JE2 and human complexes, at the top. Human complexes were then further organised by relatedness to CC8. (a-b) Arrows represent annotated genes, with colours representing key functional annotations (shared legend, right).

### SauTI1 encodes Wadjet-like condensin genes

One of the genes located on the SauTI1 is SAUSA300_1792. The NTML mutant of this gene demonstrates high frequency transfer (Supplementary Table 1), primarily through increased plasmid receipt (Supplementary Figure 3). SAUSA300_1792 has predicted homology to the condensin-family genes. Condensins are protein complexes that extrude DNA loops through their ring-like structure, folding and organising the chromosome during replication (27). Some bacteria harbour additional condensin-like complexes that recognise and digest the smaller DNA loops of plasmids; the classic example is Wadjet (19). SAUSA300_1792 shows specific homology to the structural maintenance of chromosome (*smc)* subunit gene, and we therefore designate it *smc2* here. It is also flanked by two putative exonuclease genes, *cbf1* and SAUSA300_1793, which may function to cleave recognised plasmids (Fig. 4a). This therefore provides potential homologues of JetC and JetD, the respective SMC and nuclease components of Wadjet (18). Although a knockout mutant of SAUSA300_1793 could not be obtained, the genotypes of *smc2*::Tn and *cbf1*::Tn were confirmed by sequencing. These mutants demonstrated extremely high DRP ratios across three independent repeats, suggesting that they may indeed interact with *scpB* to act as plasmid barrier genes (Fig. 4a). The slightly lower ratio of *cbf1*::Tn may be due to there being two nucleases, therefore knocking out one may be compensated by the other, resulting in a reduced phenotype in our assay. KanS DRPs had a significantly higher concentration compared to KanR DRPs in the *smc2* mutant, suggesting enhanced plasmid transfer (Fig. 4c). Furthermore, the *smc2* mutant was more susceptible to φ80α infection compared to the control strains (Fig. 4d), and its lysate contained significantly higher infectious φ80α as confirmed by plaque formation assay on RN4220 strain (Fig. 4e), which is more susceptible to phage infection compared to JE2 (Supplementary Fig. 7c). Together, these findings suggest that *smc2* functions as a key barrier to bacteriophage-mediated plasmid transfer in *S. aureus*.

**Figure 4.**
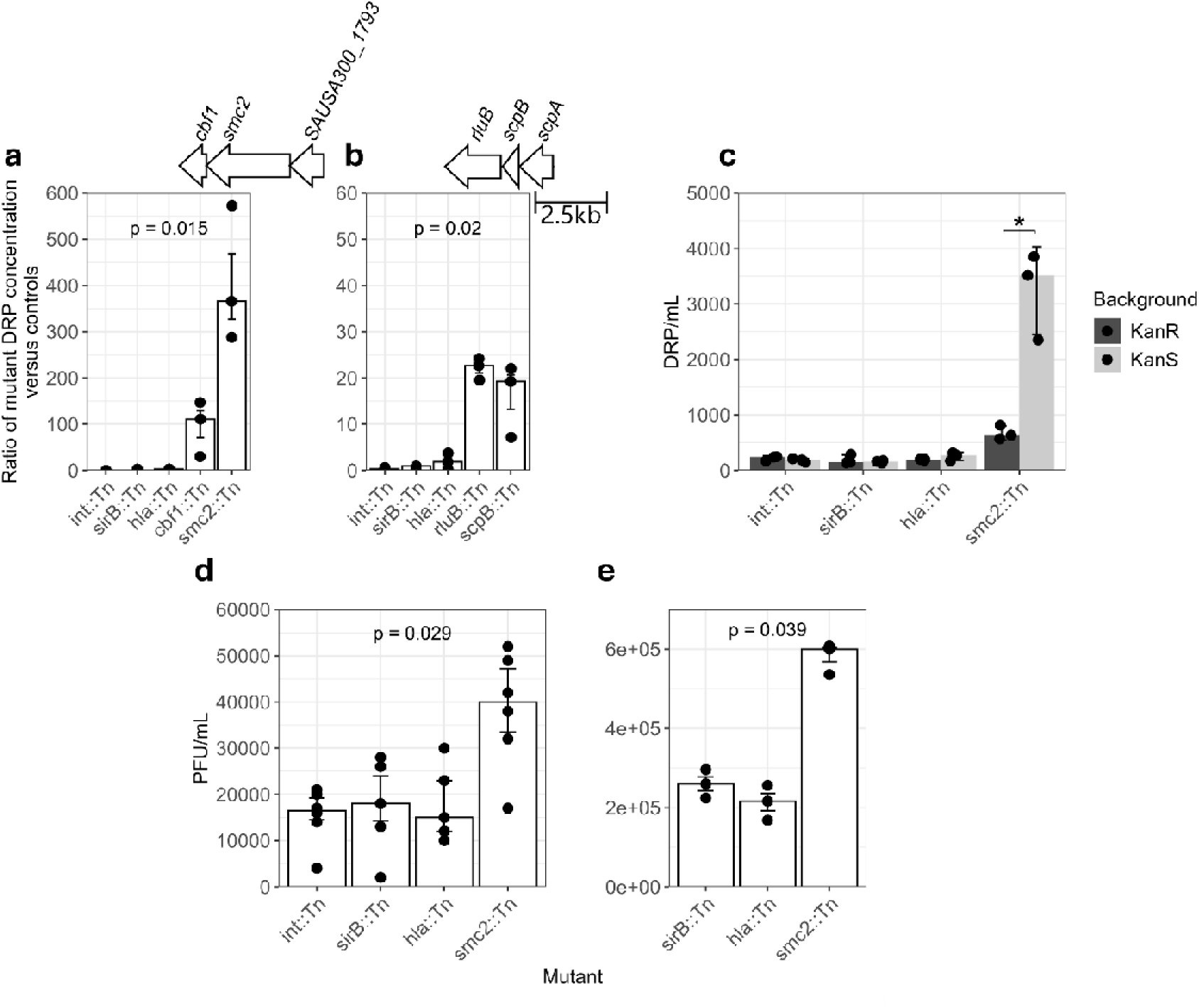
Mutants of *smc2* receive plasmid markers at high frequency and demonstrate increased phage sensitivity. (a) COGTRA analysis of co-cultures containing either control, *cbf1* or *smc2* mutants. (b) COGTRA analysis of co-cultures containing either control, *rluB* or *scpB* mutants. (a-b) Points represent the ratio of each replicate’s DRP concentration divided by the mean average control mutant DRP concentration, with bar values representing the median (n=3, per mutant). Error bars are the IQR. Statistical significance was determined by Kruskal-Wallis one-way analysis of variance. P-values are indicated. The operons are indicated by the arrows at the top of the figure; each gene is indicated by an arrow, which is laid over the bar of its respective mutant. (c) COGTRA assays of pT181 plasmid and chromosomal gene in *smc2* and control mutants showing direction of gene transfer. Kanamycin sensitive (KanS) DRPs are in the mutant background, representing transfer of the plasmid, whereas kanamycin resistant (KanR) DRPs are in the fixed strain background, representing transfer of the chromosomal gene. The mean of n=3 independent experiments is shown with SD. Statistical significance was determined by two-tailed Student’s t-test. *P<0.05. (d) Phage sensitivity assay in *smc2* and control mutants. The median of n=6 independent experiments is shown with IQR. Kruskal-Wallis one-way analysis of variance. P-values are indicated. (e) Plaque formation assay using φ80α lysates from *smc2* and control mutants on *S. aureus* RN4220. The median of n=3 independent experiments is shown with IQR. Kruskal-Wallis one-way analysis of variance. P-values are indicated.

Based on previously described Wadjet systems, the SMC protein and nuclease probably do not work in isolation (18). We therefore identified NTML mutants of condensin subunits ScpA and ScpB, homologues of JetA and JetB respectively. The *scpA* and *scpB* genes reside within an operon with the pseudouridine synthase gene *rluB* (Fig. 4b). Interestingly, *scpB*::Tn has a high rate of transfer and plasmid receipt (Supplementary Fig. 7a and Supplementary Table 3), suggesting it could be involved in controlling plasmid transfer in our system. This was unexpected, as ScpAB is demonstrated to be important in the segregation of DNA during replication (28). To ensure the high transfer phenotype was not the result of compensatory mutations, we therefore sought to generate new mutants of *smc*, *scpA*, *scpB* and *rluB* by transferring the respective knockouts into JE2. Unfortunately, correct knockout mutants of *scpA* or *smc* could not be obtained. However, having confirmed the genotype of *scpB*::Tn and *rluB*::Tn, these two mutants continued to demonstrate a high transfer phenotype (Fig. 4b). In addition, *scpB* knockout negatively impacted growth rate, suggesting wildtype gene involvement in replication, and an elevated proportion of *scpB*::Tn-background DRPs despite a relative growth rate decrease (Supplementary Fig. 7b).

## Discussion

This study identifies new genetic barriers to HGT in *S. aureus*, providing mechanistic insights into how this pathogen governs AMR acquisition. HGT is a key driver of resistance and host adaptation, occurring primarily through generalised transduction via bacteriophages in *S. aureus* (29). As internalised DNA can be potentially deleterious, bacteria must tightly control HGT to maintain genomic stability. Classical R-M systems only partially explain this control (13), and although genome-wide screens in diverse species have revealed multiple anti-phage and transfer barriers, many, including CRISPR, are rare in *S. aureus* (16, 17, 30). By screening 1,920 transposon mutants from the NTML using our COGTRA assay, which models HGT frequencies during colonisation (6), we identified 32 candidate genes that inhibit phage-mediated gene transfer in *S. aureus*. Many of these genes clustered within two chromosomal regions, designated here as SauTI1 and SauTI2, revealing new mechanisms that shape *S. aureus* evolution.

The localisation of HGT barrier genes within SauTIs suggests these loci function as coordinated defence units, allowing multiple genes to act synergically to restrict DNA transfer. Similar clustering of defence systems such as R-M and AbiE has been described in Bacillales and Enterobacterales, where co-occurring systems within defence islands (e.g., Zorya II and Druantia III in *E. coli*) collectively enhance phage resistance (31). This genomic organisation may facilitate co-regulation and mobilisation of barrier systems, conferring a fitness advantage to recipient strains. Indeed, homologs of SauTI genes were identified across diverse *S. aureus* lineages, while 11 candidate barriers within SauTI1 appear conserved within specific clonal complexes, implying lineage-specific regulation (Fig. 3). Elevated transfer of *ermB* in SauTI1 mutants (Supplementary Fig. 5a) suggests that these regions may also retain intrinsic mobilisation potential, consistent with observations in other bacterial species where barrier gene clusters are frequently associated with mobilome elements (32). Lateral transduction, which mediates efficient transfer extending hundreds of kilobases from prophage loci (33), is unlikely to explain the observed clustering, as candidate genes in SauTIs are located away from both φ2 and φ3 prophages, and φ80α does not integrate during COGTRA (21). Moreover, selected SauTI1 mutants were incapable of HGT in the absence of φ80α, confirming that the observed transfer occurs specifically via generalised transduction. The finding that SauTI mutants can influence transfer in both donor and recipient suggests that these loci may also modulate phage transduction dynamics in addition to restricting incoming DNA.

Structural Maintenance of Chromosomes (SMC) proteins are bacterial condensins essential for chromosome organisation and segregation (28, 34). Within SauTI1, we identified a condensin-like gene, *smc2* (SAUSA300_1792), and its adjacent nuclease *cbf1* as critical components of a previously unrecognised DNA restriction module that modulate phage-mediated gene transfer (Fig. 4a). Interruption of *smc2* or *cbf1* significantly increased DRP ratios, while *smc2* interruption simultaneously enhanced susceptibility to φ80α infection and plasmid transfer. The SMC2–Cbf1 system resembles the Wadjet complex described in *Bacillus cereus*, in which an SMC-like condensin (JetC) and associated nuclease (JetD) restrict the entry of circular DNA (16). Future work will be required to delineate the biochemical mechanisms underlying SMC2–Cbf1-mediated restriction and determine how they interact with phage replication or packaging. Investigating the distribution and activity of this locus across clinical isolates could also clarify its contribution to limiting the spread of plasmid-borne resistance determinants, such as resistance genes against the highly clinically important vancomycin or linezolid, which are typically transferred by conjugation (35, 36).

During this work, another screen of the *S. aureus* genome using the same mutant library was reported (37). Interestingly, *serA* was the only locus identified in both studies. Bowring, Su (37) employed temperate phage φ11 to identify loci with high transfer rates among induced mutants, an approach that primarily detects loci transferred efficiently via specialised or lateral transduction due to proximity to the φ11 integration site. Thus, the differing results of the two screens are not contradictory but reflect the distinct mechanisms each approach investigates: generalised transduction versus specialised and lateral transduction. The phenotypic screen used here was highly sensitive in detecting high-transfer mutants (HTMs), suggesting that most barrier genes were identifiable by the COGTRA assay. Over 70% of mutants showed reduced DRP ratios upon replication, and as the HTM threshold was set at the 95th percentile, a more permissive cutoff may reveal additional candidates. Mutants approaching the current threshold may therefore warrant further investigation. The identification of SauTI loci highlights genomic hotspots that may harbour additional transfer barriers or regulatory components of transduction activation pathways. Because our screen employed a fixed JE2 strain sharing the same genetic background as the NTML, it was not designed to detect systems that restrict inter-lineage or interspecies transfer. Extending this approach to genetically and ecologically diverse isolates will be critical to elucidate how SauTI activity varies across lineages, hosts, and environments, and to define its role in shaping the in vivo dynamics of gene transfer. Testing under different growth and environmental conditions, including antibiotic exposure and nutrient limitation, may further uncover additional HGT barriers in *S. aureus* (5).

In summary, this study uncovers new loci (SauTI) in *S. aureus* that form clustered genetic defence regions that restrict bacteriophage-mediated HGT in *S. aureus*. By limiting DNA exchange, these systems may contribute to the restricted inter-lineage transfer of large resistance elements such as SCC*mec*, thereby influencing the emergence and persistence of distinct MRSA lineages (5, 11). Such intrinsic genetic barriers may also constrain the dissemination of resistance, virulence, and host-adaptive genes across human and animal *S. aureus* populations (38). The identification of SauTI genes, particularly the SMC2–Cbf1 modulators of plasmid transfer, highlight their potential as biotechnological tools for genetic manipulation of *S. aureus*, offering new opportunities to study and control the spread of AMR.

## Materials and methods

### BACTERIAL STRAINS, PHAGES, AND CULTURE CONDITIONS

Bacterial strains and phages used in this study are listed in Supplementary Tables 1 and 4. The 1,920 strains of the Nebraska Transposon Mutant Library (NTML), constructed in JE2 background, were obtained from the Network on Antimicrobial Resistance in *Staphylococcus aureus* (NARSA) (20). The details of NTML mutants and their screening results are shown in Supplementary Table 1. NTML mutants indicated by BEI-prefix were obtained directly from BEI Resources, to verify genotype. The only phage used for generalized transduction was φ80α, propagated on RN4220 (39, 40).

*S. aureus* was routinely cultured at 37 °C either in brain heart infusion broth (BHIB) (Sigma-Aldrich, Dorset, UK) with shaking at 90 rpm, or on brain heart infusion agar (BHIA) unless indicated otherwise. Where appropriate, antibiotic selection was performed with erythromycin (10 μg/mL), tetracycline (10 μg/mL) and kanamycin (50 μg/mL).

### CONSTRUCTION OF SUBSTITUTION MUTANTS

To construct a fixed strain with a differing antibiotic-resistance profile for COGTRA screening, the chromosomal *ermB* gene in the φ2 integrase mutant NE201 (φ2 *int*::Tn) was replaced with the kanamycin resistance gene *aphA*-3 through allelic exchange using pKAN plasmid (41). The tetracycline-resistance plasmid pT181 was subsequently introduced from COL by transduction, generating strain NE201KT. NE201KT is erythromycin-sensitive, kanamycin-resistant, and tetracycline-resistant (42).

Clean mutants in JE2 background were generated by transduction using φ80α as previously described (43). Mutants were selected by erythromycin to ensure the transfer of the entire respective transposon insertion.

### LYSATE PROPHAGE ENUMERATION

Phage titres were determined using RN4220 as an indicator strain. Lysates were serially diluted in phage buffer (1 mM MgSO_4_, 4 mM CaCl_2_, 50 mM Tris-HCl pH 7.8, 6 g/L gelatine). For each dilution, 100 μL were combined with 400 μL of RN4220 cells in log phase (OD_600_=1) and 30 μL of 1M CaCl_2_ and incubated for 15 minutes at room temperature. These were then mixed with 7 mL phage top agar (3 g/L yeast extract, 100 mM NaCl, 3 g/L casamino acids, 3.3 g/L agar) and poured onto plates containing phage bottom agar (3 g/L yeast extract, 100 mM NaCl, 3 g/L casamino acids, 10 g/L agar). Plates were then incubated overnight, and the number of plaques on the lowest non-confluent dilution plates were counted to determine the PFU/mL. To determine the PFU in bacterial cultures and co-cultures, supernatants were collected by centrifuging 1 mL of culture (18,407 g for 3 minutes) and filter sterilized (0.2 μm) before titration.

### CO-CULTURE GENE TRANSFER RATE ASSAY (COGTRA)

COGTRA assays were performed using BHIB pre-cultures of the NE201KT fixed strain and the respective NTML mutant (1.2×10^9^ CFU/mL each) following 24-hour incubation. The pre-cultures were inoculated into 5 mL BHIB containing 20 mM CaCl_2_ to yield 1.2×10^6^ CFU/mL of each strain, followed by addition of 0.5–1×10^6^ PFU/mL φ80α. Co-cultures were incubated for 24 hours. To count DRPs, 250 µL of co-culture was plated onto BHIA containing erythromycin and tetracycline and incubated for 48 h. Unless stated otherwise, the detection limit was 4 DRP/mL. Simultaneously, each strain’s CFU was determined by plating on selective BHIA (erythromycin for NTML mutants and tetracycline for the fixed strain). Co-cultures yielding <2×10^8^ CFU/mL of either strain after 24 hours were excluded. Three control strains were included in each experiment: NE327 (φ3 *int*::Tn), NE675 (*sirB*::Tn), and NE1354 (*hla*::Tn). These loci are well characterised and not expected to influence φ80α-mediated HGT. The φ3 integrase mutant served as a partial control for φ3-mediated transfer, while *sirB* and *hla* are dispensable in iron-rich BHI (44, 45). The 1,917 non-control NTML mutants were screened following this assay (Fig1a). Mutants were tested in NTML numerical order, providing effective randomisation as NTML numbers do not correlate with chromosomal position. DRP concentrations for each NTML mutant were expressed as a ratio (R) of the mutant co-culture DRP count (D) to the mean DRP count of the three control co-cultures (µ_control_), as shown in Equation 1.

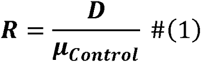

Experiments in which two or more of the control co-cultures had a DRP concentration below the limit of detection (4 DRP/mL) were rejected and repeated.

### PROPHAGE GENE MUTANT COGTRA ANALYSIS

Mutants of prophage genes predicted to be non-essential for prophage replication were analysed using the COGTRA assay in the presence and absence of φ80α. The φ3 virulence genes *scn* and *chs* were included as they are unlikely to influence replication. φ3 *dut* and the φ2 genes SAUSA300_1417 and SAUSA300_1422 have been shown to be dispensable for replication (46, 47), and the uncharacterised φ2 gene SAUSA300_1432 was also tested. All co-culture conditions were performed in triplicate.

### MULTIPLICATE COGTRA

Multiplicate COGTRA assays were performed on non-prophage HTMs identified by the screen and a selection of 49 LTMs. LTMs were selected by ordering LTMs by knockout gene locus and choosing mutants approximately every 100,000 bp (30 mutants). LTMs proximal to the SauTI1 (< 20 kb) were oversampled (6 mutants) to confirm the SauTI1 boundaries. Regions proximally up and downstream of the two prophage 2 phage headfuls were also oversampled (13 mutants) to provide mutants that are immediately adjacent to the phage (< 10 kb), within 1 headful away (< 40 kb (33)), and within 2 headfuls away (< 80 kb) to investigate whether regions surrounding the phage are low transfer, and that lateral and specialised transduction do not occur. For each mutant, assays were repeated until at least three repeats (n ≥ 3) were obtained. Due to frequent co-culture failure (parent growth below 2×10^8^ CFU/mL) mutants were frequently repeated, producing more than three replicates, with a maximum of six. The median multiplicate COGTRA ratio was then reported for each mutant.

### SCREEN SPECIFICITY AND SENSITIVITY ANALYSIS

Singular and multiplicate COGTRA results for 76 HTMs and 49 LTMs were compared to classify mutants as true HTM positives (a), false positives (b), false LTM negatives (c) or true negatives (d). Mutants that produced fewer than three successful co-cultures were excluded from this analysis. The sensitivity and specificity of the singular COGTRA was calculated using Equation 2 and Equation 3 respectively.

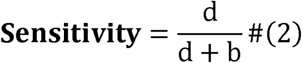

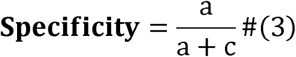

### DRP KANAMYCIN RESISTANCE ASSAY

To determine the direction of ARG transfer, DRP kanamycin resistance was tested. A sample of 25 DRP colonies were collected from each co-culture double selective BHIA plate and resuspended in 20 uL of 1X PBS. If fewer than three successful co-cultures were obtained, then the maximum number were used. Equally, if fewer than 25 DRPs were produced by a co-culture, then the maximum number was taken. Each DRP suspension (1 µL) was spotted onto triple-selective BHIA (erythromycin, tetracycline, and kanamycin) and onto double-selective BHIA (erythromycin and tetracycline) as a growth control. Plates were incubated for 24 hours. Spots showing confluent growth on both plates were classified as kanamycin-resistant (KanR), indicating the fixed-strain background; spots showing growth only on the double-selective plate were classified as kanamycin-sensitive (KanS), indicating the mutant background. The proportion of KanS and KanR DRPs in each co-culture was used to derive the DRP concentration of each background by multiplying by the total DRP count. These values were normalised by dividing by the mean DRP concentration of the corresponding control mutants, generating standardised DRP ratios.

### RELATIVE FITNESS ASSAY

The fixed strain and the respective mutant were combined in BHIB containing 20 mM CaCl_2_ at 1.2×10^6^ CFU/mL for each strain. Co-cultures were then divided into separate tubes, one per time point per co-culture, and incubated with shaking. At each time point, the appropriate co-culture tubes were removed, serially diluted in 1X PBS and plated onto BHIA containing erythromycin and tetracycline to enumerate to the CFU/mL of mutant and fixed strains. One co-culture containing fixed strain and *scpB*::Tn was excluded from further analysis as both strains failed to grow.

### GENETIC VALIDATION OF CORRECT TRANSPOSON GENE KNOCKOUT

Genotypic validation was performed for the fixed strain, control mutants, mutants of analysed beyond the initial screen, and strains produced in this study were genotypically validated. Genomic DNA was extracted from overnight cultures using the PurElute Bacterial Genomic Kit (Edge Biosystems), except for strains JW21, JW23, NE136, NE294, NE558 which were extracted using the Monarch Genomic DNA Purification Kit (New England BioLabs), following the respective manufacturer’s instructions. DNA from the fixed strain and control mutants (3–90 ng/µL in 10 mM Tris) was submitted to MicrobesNG for analysis. The remaining strains were sequenced using Illumina MiSeq. Sequence data are deposited under SRA accession PRJNA1344778 (details in Supplementary Table 4).

Reads were assembled *de novo* using Shovill (github.com/tseemann/shovill) and annotated using Prokka (github.com/tseemann/prokka) (48). Corresponding transposon resistance gene (*ermB* for all NTML mutants, with the exception of *aphA*-3 for the fixed strain) were identified by BLAST (version 2.13.0) (49). The insertion site was determined by aligning the corresponding contig to the JE2 reference genome (CP020619.1) using BLAST+ and visualised with the Artemis Comparison Tool (version 18.2.0, ACT) to locate disruption of sequence homology. (50).

### PREDICTION OF CONDENSIN-LIKE FUNCTION

Genes with predicted condensin-like functions were identified using TIGRFAM, a database using experimental as well as a range of putative evidence sources to predict the functional role of genes (51).

### SAUTI1 POPULATION DISTRIBUTION AND CONSERVATION

Twenty-two complete, well-annotated genomes representing diverse CC types were selected from Table 1 of McCarthy, Witney (52), plus JE2 (Supplementary Table 4). Multiple sequence alignments were performed using MAFFT (version 7.520), configured to automatically select an appropriate alignment strategy (53). Jalview was used to visualise and identify homologous SauTI1 loci in each sequence (54). SauTI1 loci GenBank files were obtained from NCBI (55), and visualised using Clinker (default settings) (56); JKD6009 was excluded due to lack of annotation. Sequences were sorted into CCs, organised by primary host organism (human, bovine and swine), with CC8, and therefore JE2 and human complexes, at the top. Human complexes were then further organised by relatedness to CC8, as determined by Figure 1, Baede, Gupta (57, p. 1169). CC239 was positioned beneath CC8 due to its known CC8 origin (58).

### STATISTICAL ANALYSIS & DATA VISUALISATION

The statistical analyses were performed using R as detailed in the figure legends (59). Data visualisation was performed in R using the ggplot2 package (ggplot2.tidyverse.org), unless otherwise specified (60).

To analyse whether HTM knockout genes were randomly distributed across the chromosome, 1,000 datasets were generated which randomly shuffled the DRP concentration ratios and loci values of mutants from the original dataset. Cumulative distributions of HTM genes along the chromosome for both the original and random data sets were then generated; each HTM represents a cumulative increase, therefore loci with steeper gradients represent regions with an increased concentration of HTM genes. Two-tailed Kolmogorov–Smirnov statistical comparison was then performed on the measurement difference: the mean average p-value over the 1,000 permutations is reported.

## Acknowledgements

This work was supported by Medical Research Council (grant no. 2434040 to J.W.) to MRC-LID at City St George’s, University of London and London School of Hygiene and Tropical Medicine. We are grateful to Kenneth Laing for the assistance with sequencing, Alexandra Sharpe, Arya Gupta for method development, Quentin Leclerc and Alastair Clements for the helpful discussion, and Maxmillian Wallat for help with bioinformatics.

## Contributions

J.W. performed the library screen, microbiology, statistical analyses, genomics and bioinformatics. M.M. and S.R.M. performed analysis of *smc2* mutant. M.W. and A.A.W. performed bioinformatic analysis. G.M.K. advised on statistical analyses and interpretation of results. G.M.K. and J.A.L. supervised the study including study design and data analysis. J.W. wrote the initial draft, and all authors reviewed and approved the final version.

## Ethics and consent

Ethical approval and consent were not required.

