## Supplementary Figures 1-7 and Supplementary Table 2 for "Barriers to Antimicrobial Resistance Gene Exchange in Methicillin Resistant *Staphylococcus aureus* Cluster into Transfer Islands"

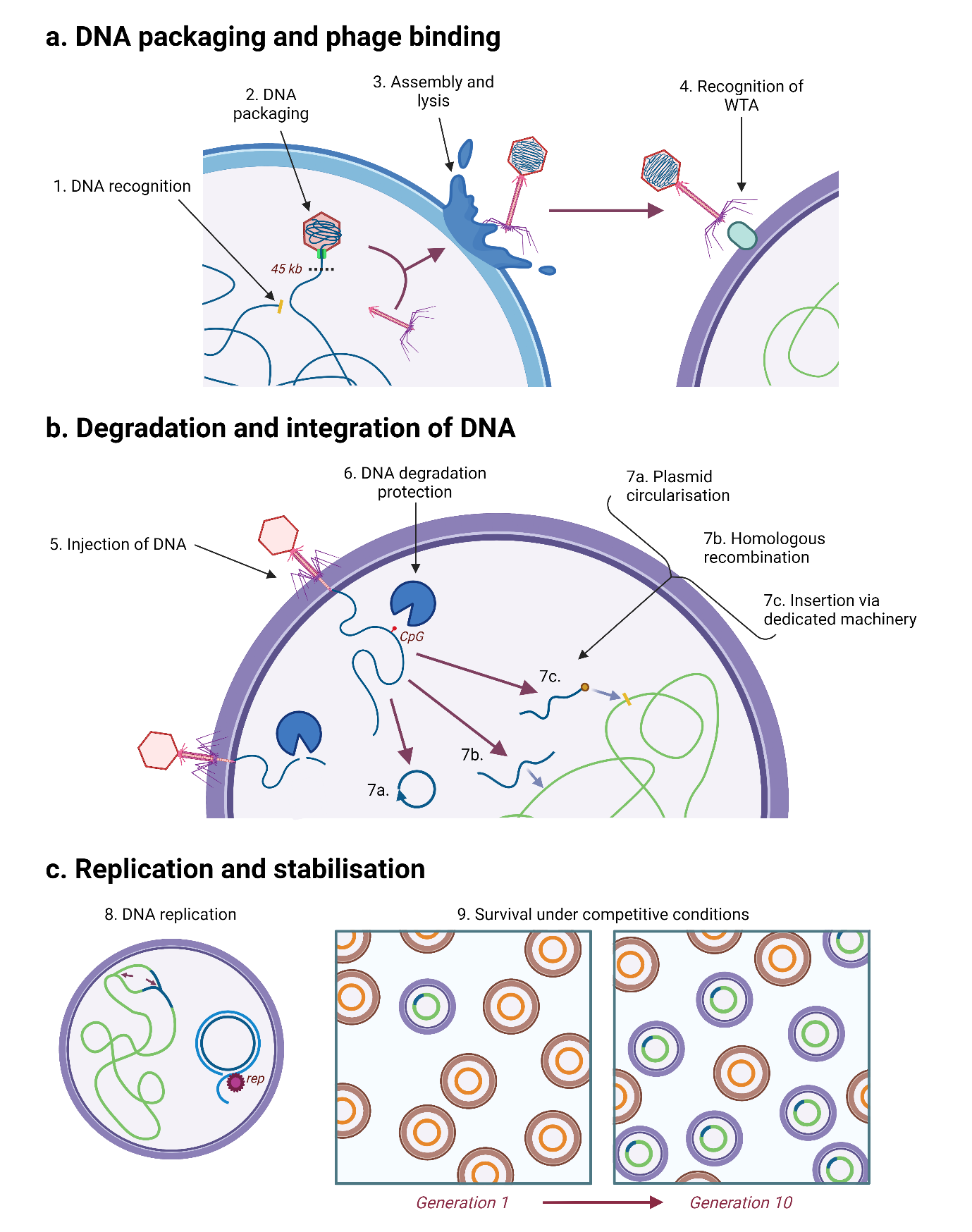

**Supplementary Figure 1. Steps to successful generalised transduction.** Bacterial DNA is initially recognised by phage terminase machinery (1), which then packages the DNA into the phage capsid until headful capacity is reached (2). Mature transducing particles are assembled and released from the donor cell via lysis (3). Transducing particles recognise and bind the wall teichoic acids (WTA) of a new bacterium (4). This triggers the injection of donor DNA into the recipient cell (5), which must then evade degradation by the recipient’s nuclease machinery, for example target sites become methylated (CpG) to evade restriction modification system recognition (6). If the donor DNA is a plasmid, circularisation will often occur (7a). Chromosomal DNA, however, must integrate into the genome via homologous recombination (7b), an inefficient process; some mobile genetic elements, such as *Staphylococcus aureus* pathogenicity islands and transposons, have dedicated machinery that facilitates efficient insertion, sometimes into dedicated insertion sites (7c). Donor DNA that has inserted into the recipient chromosome is replicated with it, however plasmid DNA must autonomously replicate using its own replicative machinery (*rep*) (8). For DNA to remain stable within the population it must not negatively impact cell fitness, otherwise it may be lost or cause the host bacteria to be outcompeted (9).

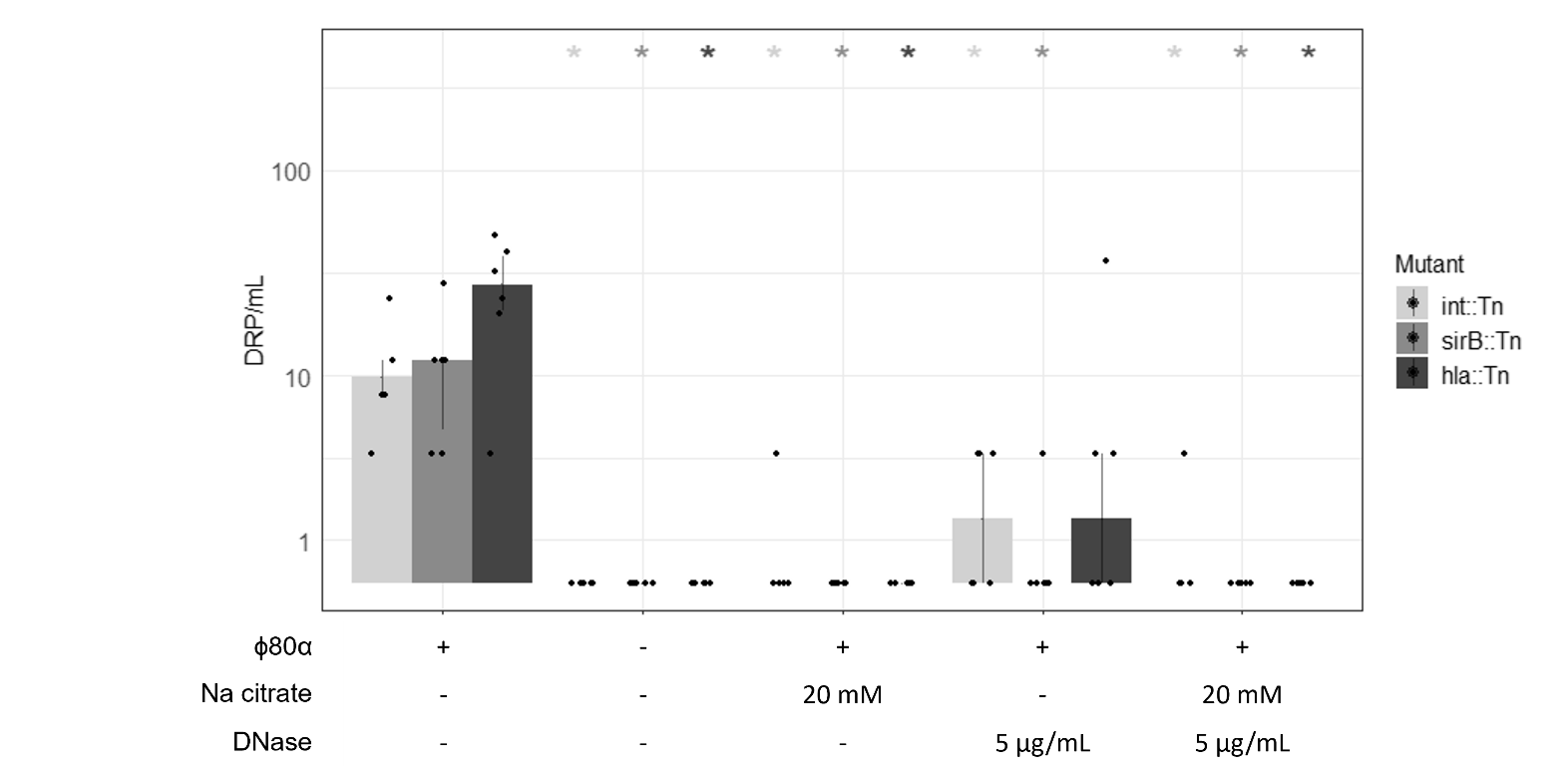

**Supplementary Figure 2. Transformation is unlikely to contribute to DRP generation in COGTRA experiments.** Control mutant COGTRA in the presence or absence of ɸ80α, 20 mM Na citrate, and/or 5 µg/mL DNase. Points represent the DRP concentration of each replicate (n = 6) after 24 hours, with bar values representing the median. Error bars are the IQR. Each mutant condition was compared to their respective mutant control, containing ɸ80α and no Na citrate or DNase, via Wilcoxon rank-sum test with Bonferroni correction. Asterisks denote adjusted p-values (no asterisk > 0.05, * < 0.05, ** < 0.01, *** < 0.001), with colour denoting the NTML control mutant.

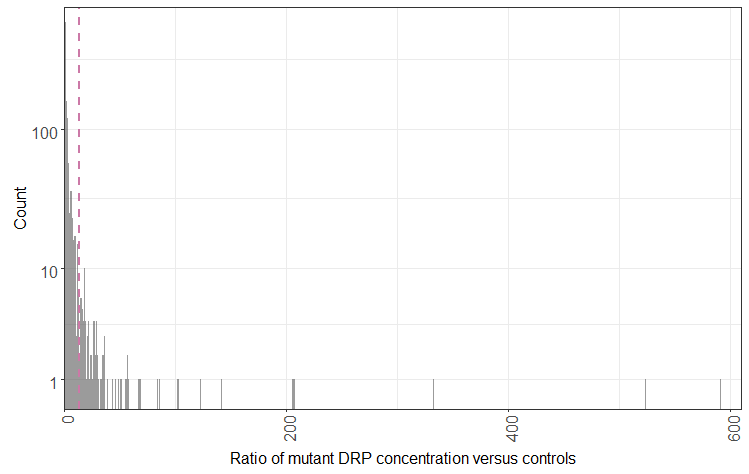

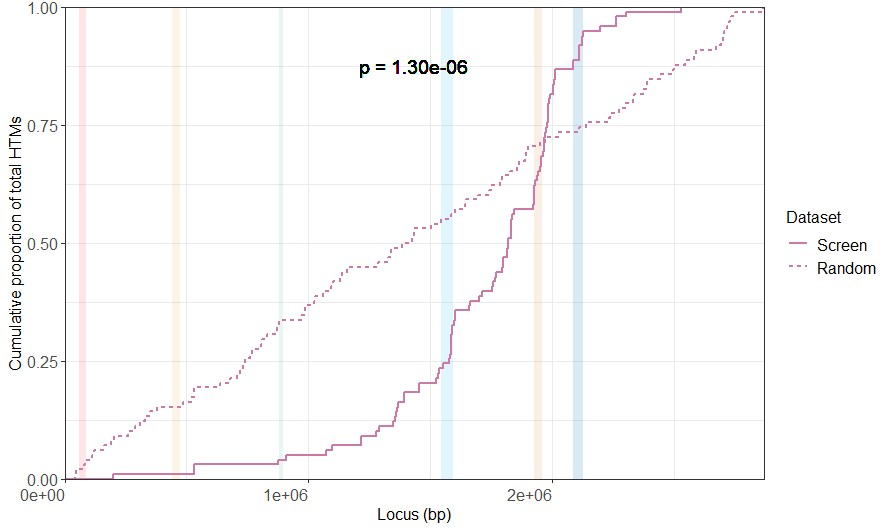

**a.**

**b.**

**Supplementary Figure 3. HTM genes are non-randomly distributed along the *S. aureus* chromosome.** (a) Histogram of mutant DRP concentrations vs controls in the single COGTRA screen. Binwidths are a ratio of 1**.** The 95^th^ percentile (ratio = 13) is indicated by the pink dashed line. (b) The cumulative proportion of HTM genes along the chromosome from the COGTRA screen (solid) and a single randomly shuffled distribution of HTMs dataset (dashed). The presence of a HTM knockout gene at a locus is represented by a cumulative increase. Genomic features are indicated with light coloured vertical bars (SCC*mec* = red, νSAα = yellow, SaPI5 = green, ɸ2 = light blue, νSAβ = orange, ɸ3 = dark blue). The p-value was calculated via a two-tailed Kolmogorov–Smirnov test. The SauTIs are indicated within red boxes. To ensure that this result was robust, 1000 additional random datasets were generated and compared to the original dataset. The mean average p-value of these additional permutations was 6.80x10-5.

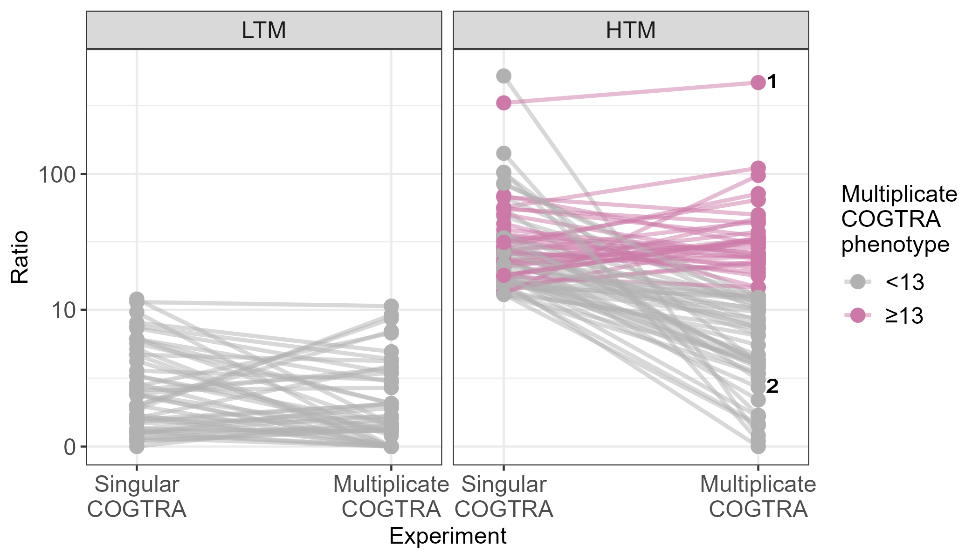

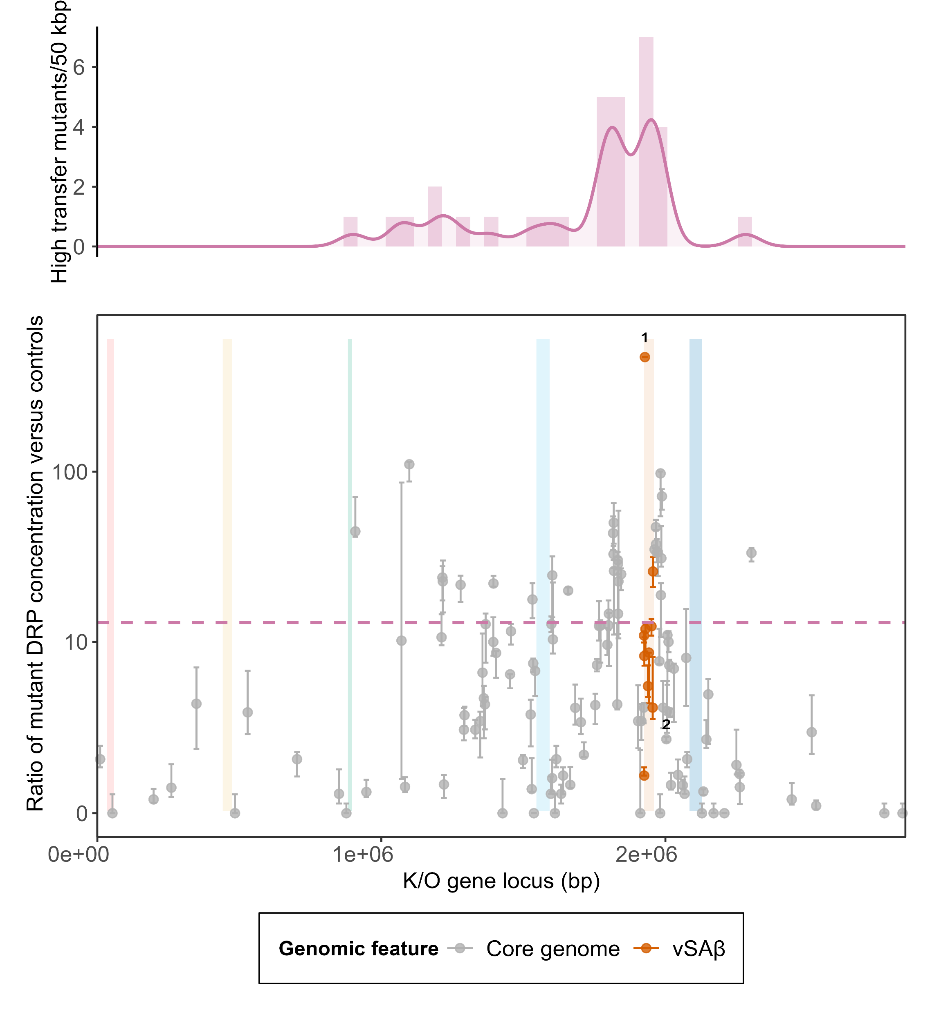

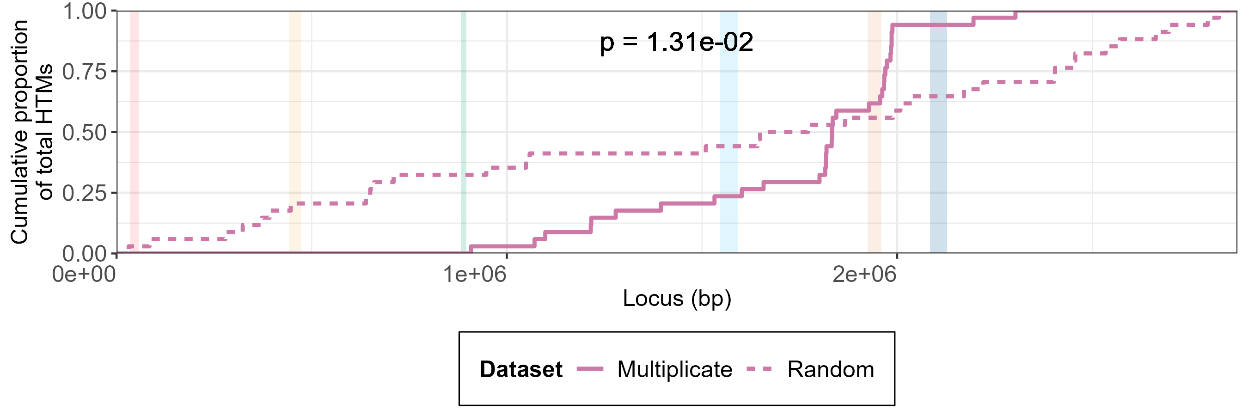

**d.**

**c.**

**b.**

**a.**

**Supplementary Figure 4. Confirmed high frequency transfer library mutants have knockout genes that are non-randomly clustered between, but not next to, prophage.** (a) The singular COGTRA ratio of 49 low transfer mutants and 78 core high transfer mutants was plotted against the corresponding multiplicate COGTRA ratio (y-axis). Ratios were calculated by dividing the DRP concentration of each co-culture by the mean DRP concentration of three control mutants (*int*::Tn, *sirB*::Tn and *hla*::Tn; control co-culture repeats: singular, n=1; multiplicate, n=3). Multiplicate COGTRA values represent the median ratio; 3≤n≤6, except SAUSA300_1747::Tn^1^ (n=1) and SAUSA300_1842::Tn^2^ (2, n=2). Lines connect the singular and multiplicate COGTRA ratios of each mutant. Colours represent whether the multiplicate COGTRA experiment median ratio ≥ 13 (legend right). (b) An overlayed histogram of confirmed HTM knockout genes per 50 kb; trend line (solid pink) plotted via kernel density estimation. (c) The DRP concentration of each mutant co-culture repeat divide by the experiment’s mean DRP concentration of control mutant co-cultures (y-axis) was plotted against the start locus of the corresponding knockout (K/O) gene (x-axis) to identify where confirmed HTM knockout genes are located on the chromosome. Points represent the median ratio of each mutant; error bars represent the IQR; 3≤n≤6, except SAUSA300_1747::Tn1 (n=1) and SAUSA300_1842::Tn2 (2, n=2). High transfer mutant threshold (≥ median ratio of 13) is represented by the pink dashed line. (d) The cumulative proportion of confirmed HTM genes along the chromosome from the multiplicate COGTRA (solid) and a single randomly shuffled distribution of HTMs dataset (dashed). The presence of an HTM knockout gene at a locus is represented by a cumulative increase. The p-value was calculated via a two-tailed Kolmogorov–Smirnov test. Genomic features are indicated with coloured vertical lines (SCCmec = red, νSAα = yellow, SaPI5 = green, ɸ2 = light blue, νSAβ = orange, SauTI1/2 = pink, ɸ3 = dark blue). To ensure that this result was robust, 1000 additional random datasets were generated and compared to the original dataset. The mean average p-value of these additional permutations was 0.016.

**Supplementary Table 2. Contingency table of singular COGTRA screen HTM-phenotype identification compared to multiplicate repeat gold standard.**

|  | | **Multiplicate COGTRA (3≤n≤6)** | |
| --- | --- | --- | --- |
|  |  | LTM | HTM |
| **Singular COGTRA (n=1)** | LTM | 49  (True Negative) | 0  (False Negative) |
|  | HTM | 44  (False Positive) | 32  (True Positive) |
|  |  | Specificity = 52.7% | Sensitivity = 100% |

Note: Abbreviations – LTM, low transfer mutant (ratio < 13); HTM, high transfer mutant (ratio ≥ 13)

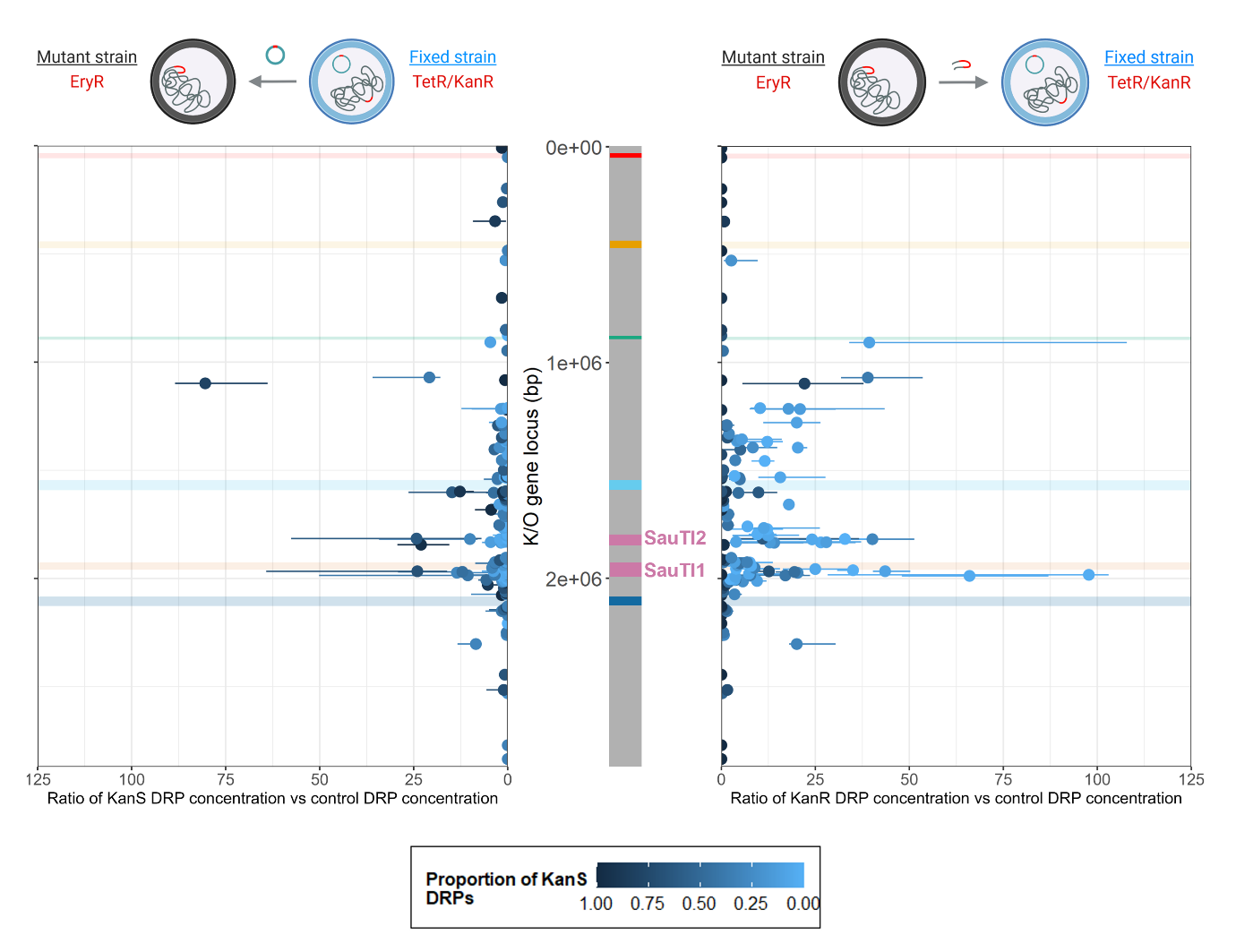

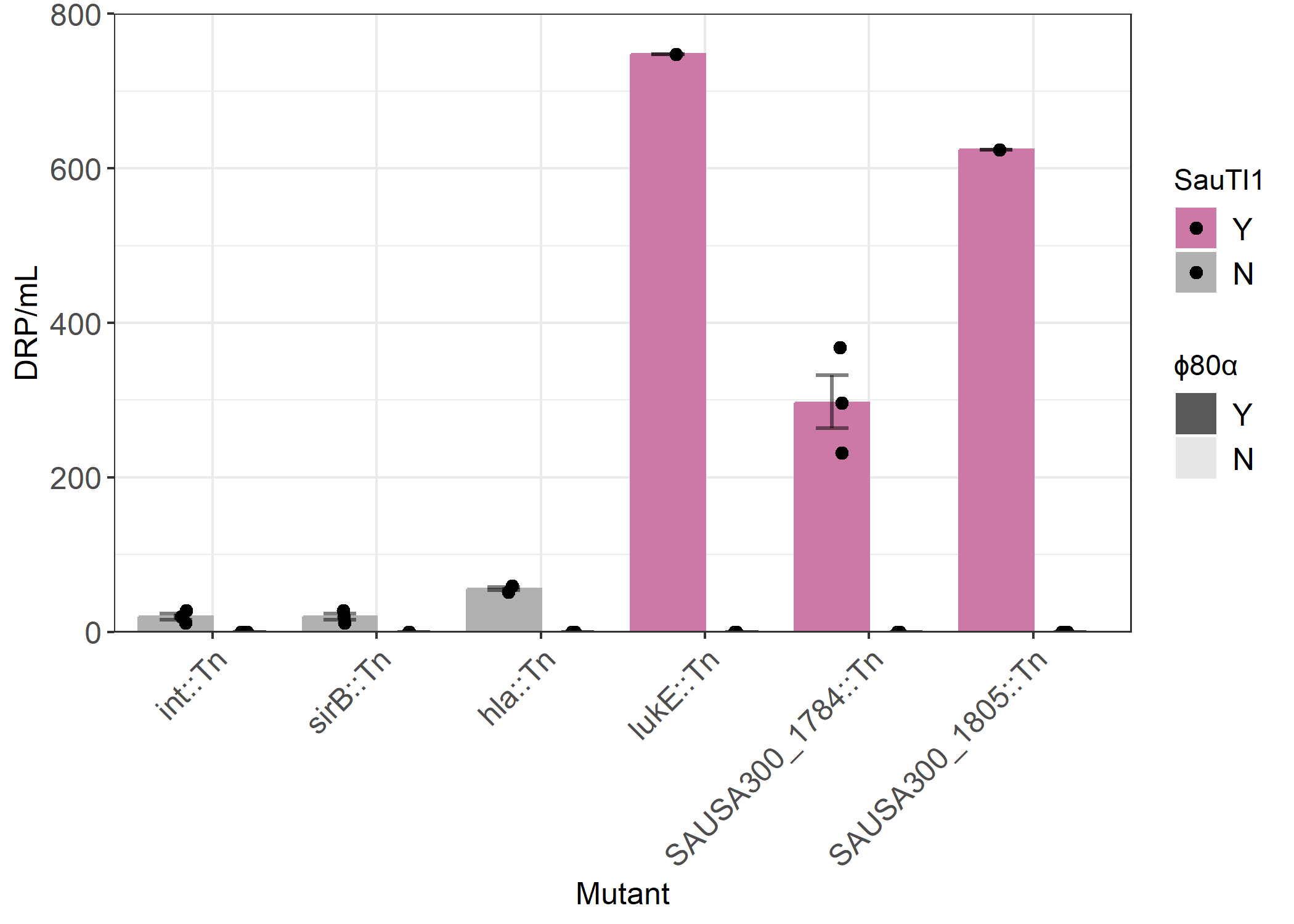

**a.**

**b.**

**Supplementary Figure 5. *Staphylococcus aureus* Transfer Island 1 (SauTI1) loci mutants show increased transfer in either direction, or both, requiring ɸ80α.** (a) The rates of (left) tetracycline resistance plasmid (TetR) acquisition and (right) erythromycin resistance gene (EryR) donation, as represented by the top graphics, were measured for 94 confirmed LTMs and 33 confirmed HTMs. The concentrations of (left) kanamycin sensitive (KanS) or (right) resistant (KanR) DRPs were divided by the mean overall DRP concentration of three control mutants (*int*::Tn, *sirB*::Tn and *hla*::Tn; control co-culture repeats: n=3) to control for experimental variation. Ratios (horizontal y-axes) were plotted against the start locus of the corresponding knockout (K/O) gene (shared vertical x-axis). Points represent the median ratio (n = 3); error bars represent the IQR; point/error bar colours represent the mean proportion of kanamycin sensitive DRPs of the sample (n ≤ 25) tested from each co-culture, which were used to calculate the KanS/KanR concentrations. Genomic features are indicated with horizontal colours (SCC*mec* = red, νSAα = yellow, SaPI5 = green, ɸ2 = light blue, νSAβ = orange, SaPI1/2 = pink, ɸ3 = dark blue). The y-axis limit was set at a ratio of 125, which excluded SAUSA300_1747::Tn (gene locus = 1927784 bp; median KanS ratio = 337.8; median KanR ratio = 131.3; n = 1). (b) COGTRA of mutants of genes located at the start (*lukE*::Tn), middle (SAUSA300_1784::Tn) and end (SAUSA300_1805::Tn) of the JE2 SauTI1, as well as 3 control mutants, were analysed via in the presence or absence of ɸ80α. Points represent the DRP concentration of each replicate (n = 3), with bar values representing the median. Bar colour represents whether the knockout gene is located on the SauTI1; bar transparency represents whether ɸ80α was present in the co-culture. Error bars are the IQR.

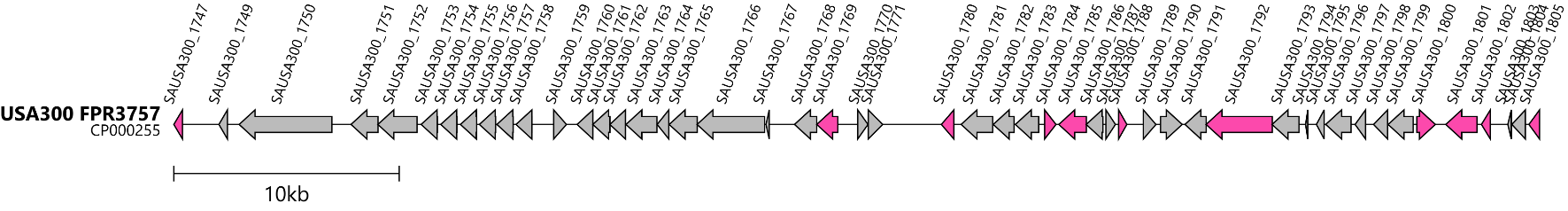

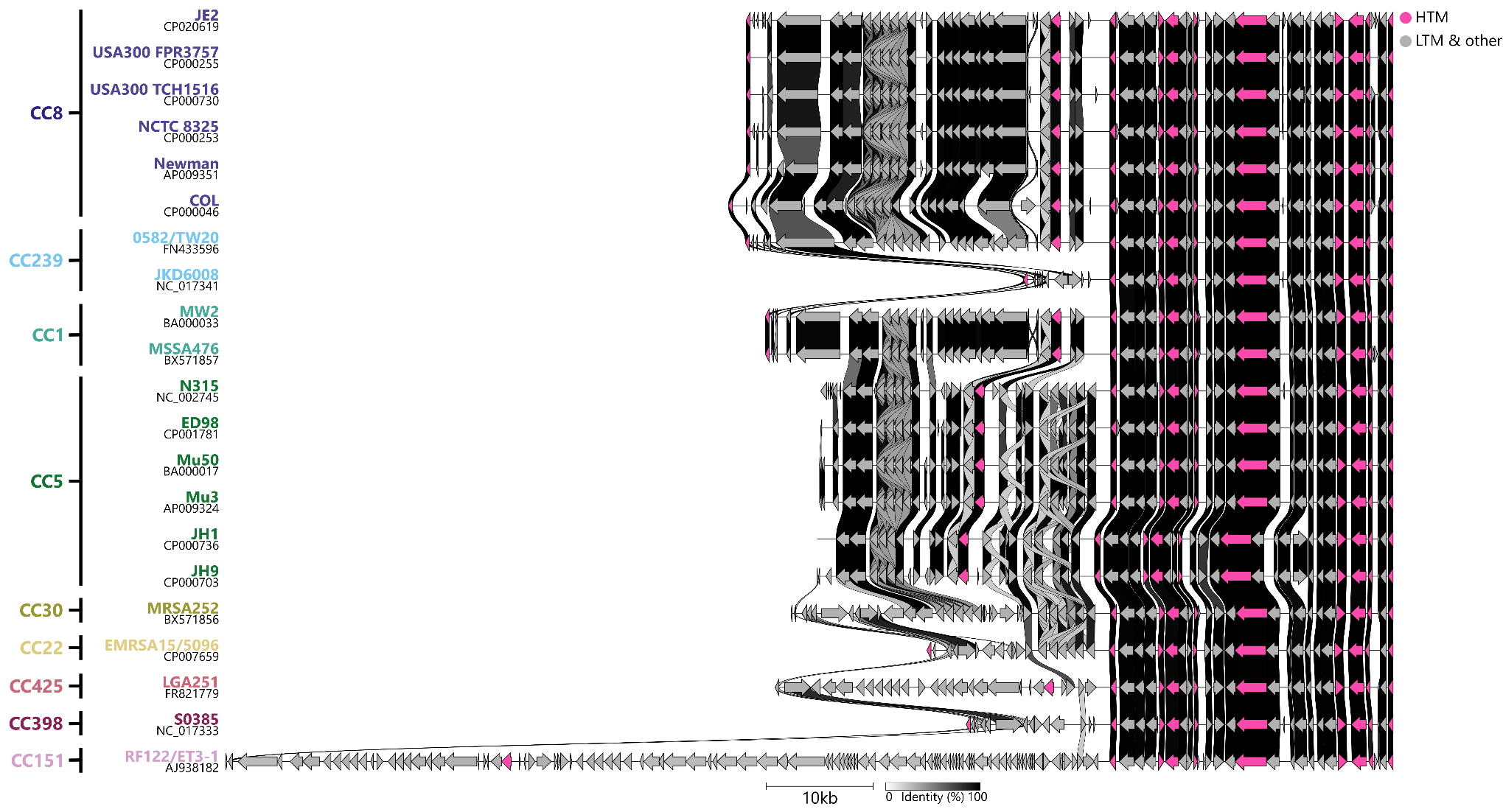

**a.**

**b.**

**Supplementary Figure 6. Gene homology between SauTI1 HTMs by clonal complex (CC).** (a) Schematic map of the USA300_FPR3757 SauTI1. The respective locus is presented at the start of each gene. (b) Comparison of the SauTI1 homologues of 21 isolates. Isolate name and accession number are left of the figure, with respective colour representing the respective CC. Connecting bands represent the degree of overall identity between the gene homologues stacked above it (identity bar, bottom). (a-b) Arrows represent annotated genes, with colours representing confirmed HTM genes (shared legend, right).

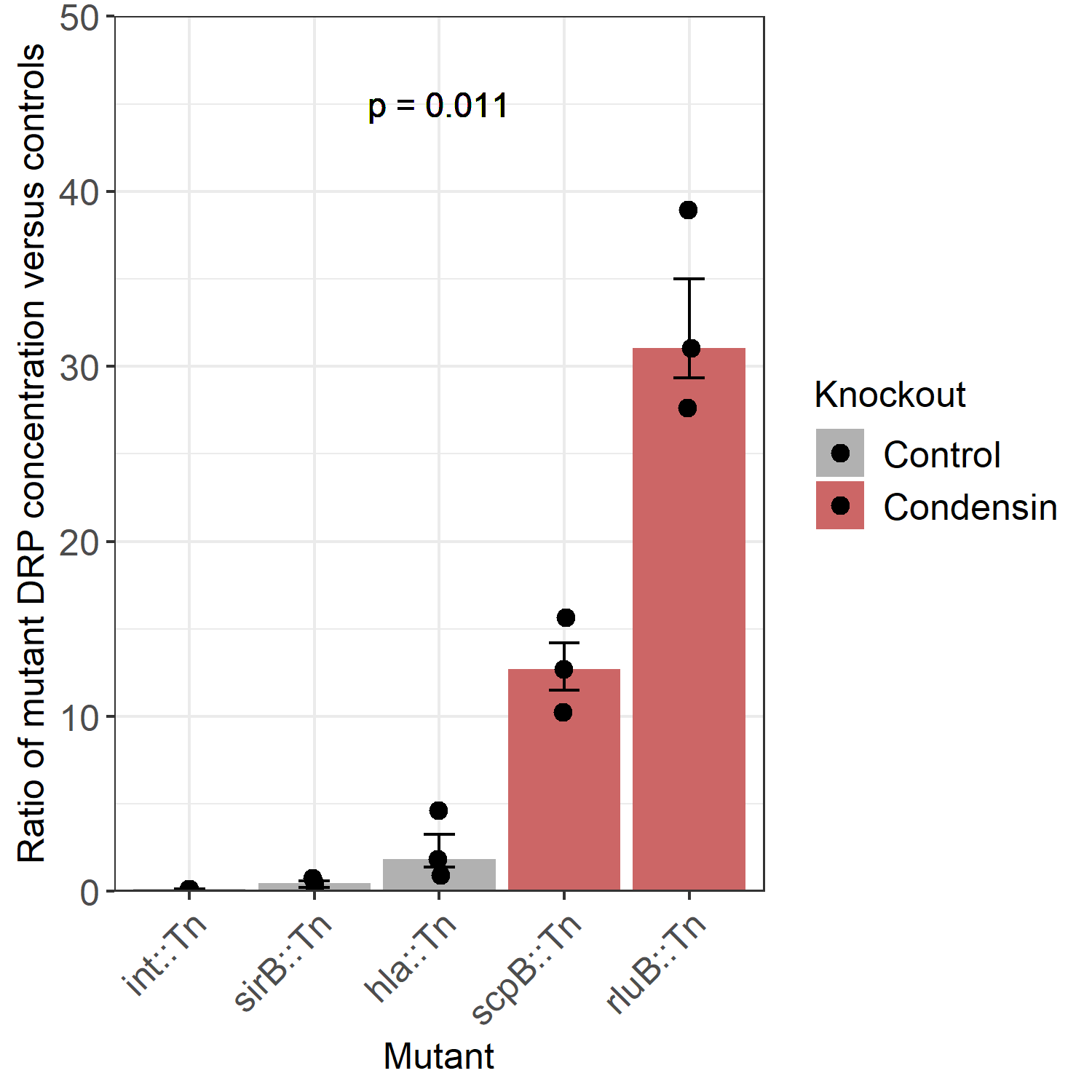

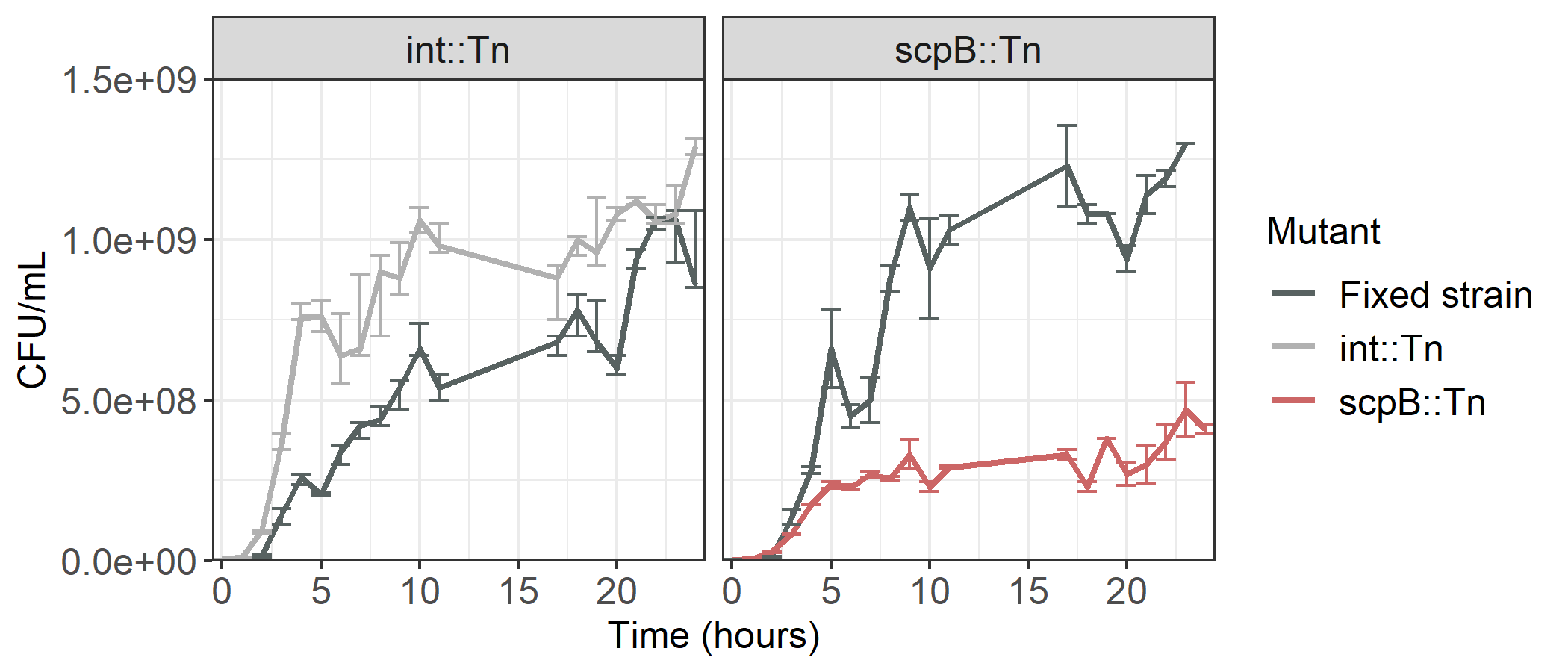

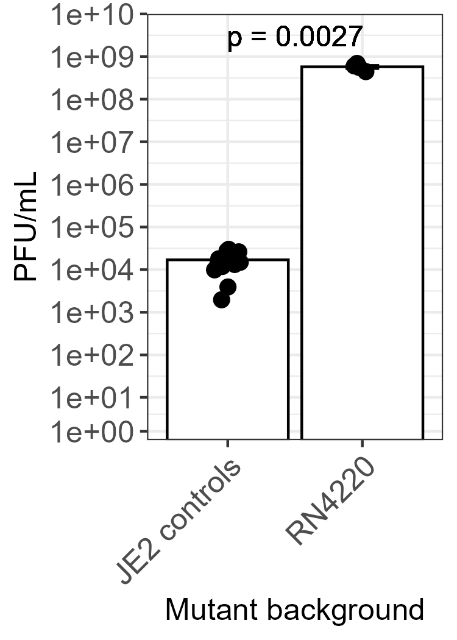

**c.**

**a.**

**b.**

**Supplementary Figure 7. Mutants of *scpB* demonstrate increased HGT despite reduced fitness.** (a) COGTRA analysis of co-cultures containing control, *scpB* or *rluB* mutants were determined. Points represent the ratio of each replicate’s DRP concentration divided by the mean average control mutant DRP concentration, with bar values representing the median (n = 3, per mutant). Error bars are the IQR. The p-value was calculated via Kruskal-Wallis one-way analysis of variance. Although *rluB* is not a condensin gene by function, it has been included due to its situation within the *scpAB* operon, and for the purposes of the figure. (b) Bacterial concentrations in co-cultures containing the fixed strain and either *int*::Tn (n = 3) or *scpB*::Tn (n = 2), in the absence of ɸ80α, were measured over 24 hours. Colour represents the respective mutant. (c) Phage sensitivity assay in JE2 control mutants (*int*::Tn, *sib*::Tn, *hla*::Tn) and RN4220. The medians of independent experiments of RN4220 (n = 4) and JE2 control mutants (n = 18) are shown with the IQR. The results of the control mutant experiments were combined (n = 6 per mutant). Statistical significance was determined by Wilcoxon rank-sum test.
